# Benchmarking porcine pancreatic ductal organoids for drug screening applications

**DOI:** 10.1101/2025.03.12.642843

**Authors:** Christos Karampelias, Kaiyuan Yang, Michael Sterr, Mireia Molina van Den Bosch, Simone Renner, Janina Fuß, Sören Franzenburg, Tatsuya Kin, Eckhard Wolf, Elisabeth Kemter, Heiko Lickert

**Affiliations:** Institute of Diabetes and Regeneration Research, Helmholtz Munich, Neuherberg, Germany; German Center for Diabetes Research (DZD), Neuherberg, Germany; School of Medicine and Health, Technische Universität München, Munich, Germany; Gene Center and Center for Innovative Medical Models (CiMM), LMU Munich, Munich, Germany; Interfaculty Center for Endocrine and Cardiovascular Disease Network Modelling and Clinical Transfer (ICONLMU), LMU Munich, Munich, Germany; Institute of Clinical Molecular Biology, Christian-Albrechts-Universität Kiel, Kiel, Germany; Clinical Islet Laboratory, University of Alberta Hospital, Edmonton, AB, Canada

**Keywords:** Pancreatic ductal organoids, porcine pancreas, chemical screen, cross-species comparison, scRNA-Seq profile

## Abstract

Primary human pancreatic ductal organoids (HPDO) have emerged as a model to study pancreas biology and disease. Yet, donor material availability, and a lack of extensive benchmarking limits the range of applications. To address this gap, we established porcine pancreatic ductal organoids (PPDO) as a system from an easily obtainable source to model pancreatic ductal/progenitor biology. We benchmarked PPDO to HPDO and primary porcine pancreas using single-cell RNA sequencing (scRNA-Seq). We observed no overt phenotypic differences in PPDO derived from distinct developmental stages, with a WNT signaling enriched population characterizing PPDO. PPDO exhibited differentiation potential towards mature ductal cells and limited potential towards endocrine lineages. We used PPDO as a platform to assess the safety of FDA-approved drugs and showed conserved toxicity of statins and α-adrenergic receptor inhibitors between PPDO and HPDO cultures. Overall, our results highlight the PPDO as a model for mammalian duct/progenitor applications.

## Introduction

The homeostatic turnover of the mammalian adult pancreas is a relatively slow process and has been postulated to rely mainly on cell proliferation^1–4^. Whether the adult human pancreas contains self-renewing and/or adult stem cells that contribute to exocrine or endocrine tissue (re)generation is still an open question. It is known that the embryonic multipotent ductal epithelium gives rise to all endocrine, exocrine and ductal cells in mammals, but *in vivo* lineage-tracing studies have shown that ductal progenitors are lineage restricted and do not contribute to other lineages in the adult pancreas^3,5–8^. Exocrine tissue maintenance and regeneration is less studied. Recent work suggested that acinar cell maintenance occurs mainly through self-replication, while some studies indicated the presence of acinar progenitor populations that contribute to tissue maintenance and might be involved in carcinogenesis^4,9^. Studies on identifying adult human pancreatic progenitors are limited, yet there are indications that postnatal ductal cells could behave as such^10–14^. Given the limitations to study human adult pancreatic ductal/progenitor biology in large scale, new and thoroughly characterized organoid models are needed to advance basic biology and regenerative medicine.

Organoids are 3D culture models initially established from tissues known to harbor adult stem cells in their epithelium (e.g. the intestine) and their properties are being standardized according to recent community guidelines^15,16^. In this regard, duct-derived pancreatic organoids from primary tissue can be used as a proxy to address questions on adult pancreatic progenitor and/or ductal biology and disease. Primary adult tissue pancreatic organoids were first generated from mouse ductal cells^17–20^. Several groups have generated human ductal organoids both from exocrine pancreatic tissue and from pluripotent stem cells derived pancreatic progenitors^13,19,21–29^. So far, pancreatic ductal organoids have mainly been used to model pancreatic cancer and to study endocrine cell differentiation to identify disease mechanisms and advance regenerative therapies. However, human pancreatic tissue for organoid generation is limited by the availability of donor material and suffers from fast degradation due to acinar enzyme mediated digestion. Additionally, stem cell-derived organoids only resemble a more embryonic progenitor phenotype. This highlights the need for new, relevant and thoroughly phenotyped and characterized pancreatic organoid models to advance our knowledge of pancreatic adult progenitor populations and pancreatic disease.

To address this technological gap, we generated and characterized in-depth porcine pancreatic ductal organoids (PPDO) from healthy and diabetic pigs^30^ across distinct developmental stages. We used PPDO to model ductal/progenitor biology dynamics *in vitro* and to study drug-induced exocrine-related pathologies (pancreatitis-pancreatic cancer). The pig serves as a large animal model to study pancreas biology with its advantages being resemblance in terms of size and metabolism to humans coupled to efficient transgenesis for disease modeling^31^. Moreover, the availability of porcine pancreata from developmental stages (embryonic, early post-natal) that are difficult to obtain from human donors, makes them an ideal surrogate to study pancreas biology and disease using organoids. In this work, we observed that PPDO were transcriptional- and protein-wise similar across ages and genotypes, yet their molecular signature changed after prolonged maintenance in culture. To evaluate their suitability for translational applications, we performed a chemical screen using PPDO to assess the safety of FDA approved drugs for pancreas diseases, translating these findings to HPDO. Overall, we provide in-depth benchmarking and application of PPDO as a model for adult pancreatic ductal/progenitor biology.

## Results

### Generation and characterization of PPDO across developmental stages

We generated PPDO from primary porcine pancreatic ductal cells across different ages (embryonic (E) and postnatal (PN)) and genotypes (wild type, *INS*-eGFP^32^, *INS*- eGFP/*INS*-C94Y^30^). This is a unique advantage using the pig model to study human-relevant biology across distinct developmental ages and with different transgenic models. Briefly, we manually picked ductal structures following pancreas digestion and cultured them with extracellular matrix (Cultrex) in human pancreatic ductal organoid (HPDO) medium^20^ (Figure 1A). Phenotypically, PPDO initially formed large cystic structures, like the HPDO derived from human pancreatic exocrine fractions. We did not observe any differences in organoid formation capacity or overall morphology between PPDO generated from embryonic compared to postnatal stages or from healthy compared to diabetic pig pancreata (Figure 1B-D). PPDO cultures could be freeze-thawed and were maintained up to passage 12, at which time point the cultures were cryopreserved due to experimental protocol ending. However, we noticed a phenotypic switch of the PPDO upon prolonged culturing, with the cystic morphology giving rise to smaller, dense cellular spheres without a discernible lumen. This phenotype appeared between passages three to five and the dense sphere structure became dominant following passage six (Figure 1B’-D’). No such morphological transformation occurred for HPDO in our hands, although the HPDO size became smaller over increasing passage number (Figure 1E-E’).

**Figure 1:**
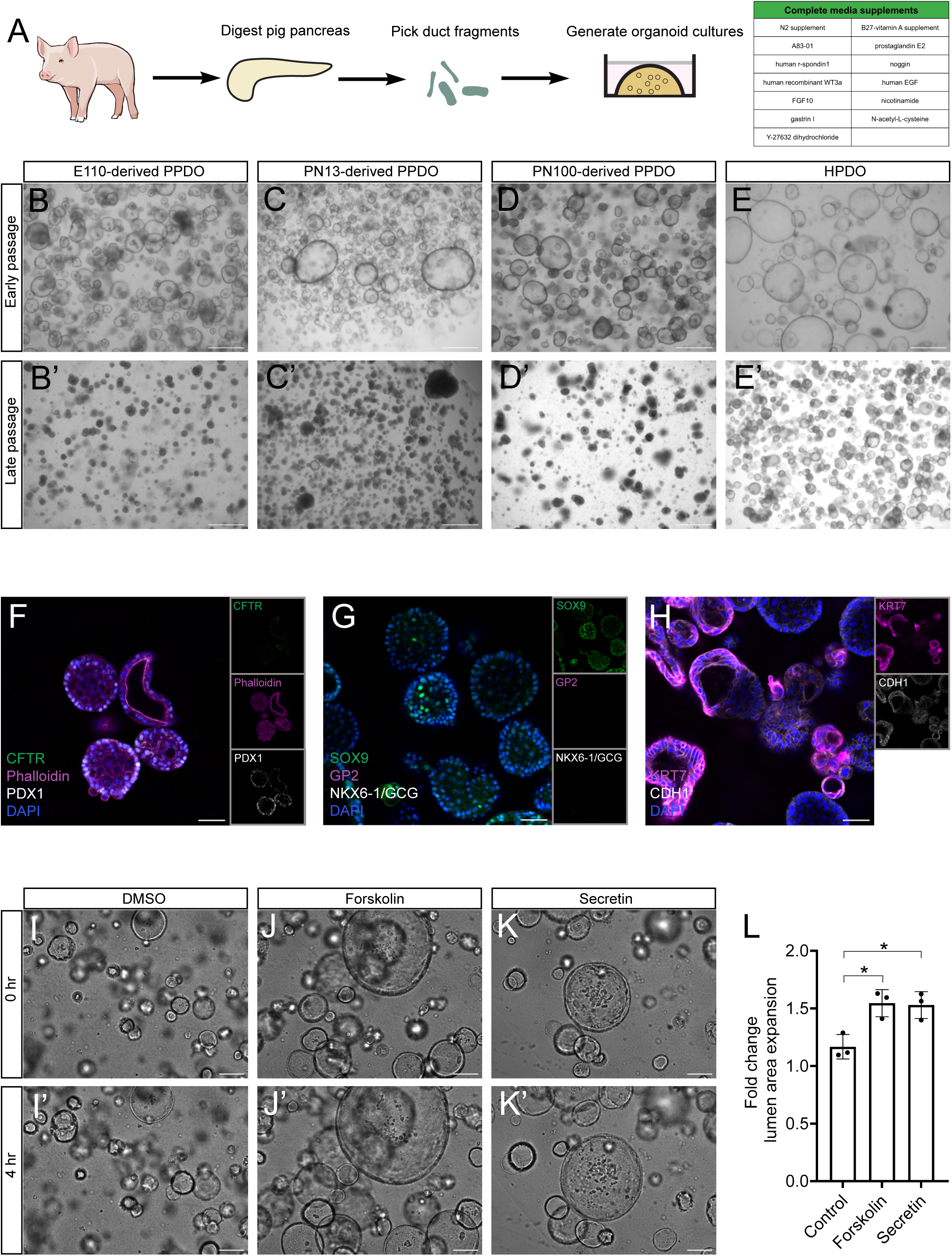
Generation and characterization of PPDO across developmental stages. (A) Schema showing the process for generating PPDO (B-E) Brightfield microscopy images of early passage PPDO and HPDO from E110- WT (B), PN13- *INS*-eGFP/INS-C94Y (C), PN100-*INS*:eGFP (D) and from a human donor (E). Scale bar 500 µm. (B’-E’) Brightfield microscopy images of late passage PPDO and HPDO from E110- WT (B’), PN13- *INS*-eGFP/*INS*-C94Y (C’), PN100-*INS*:eGFP (D’) and from a human donor (E). Scale bar:500 µm. (F-H) Single-plane confocal images of PPDO derived from PN19 pig pancreas immunostained against CFTR-Phalloidin-PDX1 (F), SOX9-GP2-NKX6-1/GCG (G), KRT7-CDH1 (H) and counterstained with DAPI. Insets show the individual channels of the merge image. Scale bar: 50 µm. (I-K) Brightfield microscopy images of early passage PPDO at the beginning of the live imaging, treated with DMSO (I), forskolin (J) and secretin (K). Scale bar 500 µm. (I’-K’) Brightfield microscopy images of early passage PPDO at the end of the live imaging, treated with DMSO (I’), forskolin (J’) and secretin (K’). Scale bar 500 µm. (L) Quantification of the lumen area expansion of PPDO following 4h of live imaging and treatments with forskolin and secretin. n=3 independent experiments with different PPDO lines. One-way ANOVA followed by Dunnett’s multiple comparisons test was used to assess significance. Padj*0.015 for DMSO vs forskolin and *0.0140 for DMSO vs secretin.

We profiled key pancreatic protein markers in PPDO, including, acinar/pancreatic progenitor and endocrine specific proteins. Immunostaining confirmed that the PPDO express ductal (CF transmembrane conductance regulator (CFTR), SRY-Box transcription factor 9 (SOX9), keratin 7 (KRT7)) and epithelial markers (Pancreatic and duodenal homeobox 1 (PDX1), cadherin 1 (CDH1)), markers but were devoid of endocrine (NK6 Homeobox 1 (NKX6-1), glucagon (GCG) and acinar cell markers (glycoprotein 2 (GP2). F-actin (phalloidin) immunostaining showed higher intensity in the luminal area of the PPDO suggesting apical-basal polarization of the epithelium (Figure 1F-H). This protein expression pattern was consistent across PPDO derived from different developmental stages (Figure S1A-I). Moreover, we assayed ductal functionality of early passage PPDO with the CFTR functionality assay, utilizing both forskolin (non-physiological) and secretin (physiological) to activate ductal cell bicarbonate excretion. If the cultures include mature, CFTR-expressing ductal cells, stimulation will lead to fluid uptake and lumen expansion. Lumen area of PPDO significantly expanded following treatment with both agonists (Figure 1I-L) and across PPDO derived from different ages, suggesting functional ductal cells in terms of bicarbonate secretion. Overall, we established and showed the ductal fate of PPDO from distinct porcine developmental stages, a fate that appeared to change upon prolonged culturing.

### Deep phenotyping of PPDO and benchmarking to HPDO by scRNA-Seq

We used scRNA-Seq to transcriptionally profile PPDO and benchmark them against the state-of-the-art HPDO cultures. We sampled two PPDO lines (PN100/PN240), which were cultured for four and seven passages respectively, to better understand the observed cell fate/morphological transformation. Similarly, we performed scRNA-Seq profiling of two HPDO lines originating from adult human pancreatic ductal cells at passage four and seven mixed in one sample. For the PPDO dataset, we identified seven cell populations/states corresponding to pancreatic ductal cells, a proliferating population, a potential progenitor population characterized by high WNT signaling, a population/state characterized by expression of basal polarity genes and two populations/states that were enriched for glycolysis relating genes and processes (Figure 2A&C). This transcriptional signature was similar between the two PPDO lines assayed, a result that showcased the similarity of PPDO cultures regardless of the developmental stage they derived from. PPDO were enriched for expression of ductal marker genes (solute carrier family 4 member 4 (*SLC4A4)*-bone morphogenetic protein receptor type 1A (*BMPR1A)*-*KRT7*) although at these later passages *SOX9* and mucin 1, cell surface associated (*MUC1*, *ENSSSCG00000006525* in pig genome) expression was relatively low and restricted to the pancreatic ductal population 2. No significant expression of other pancreatic cell type markers was observed (Figure 2B&D). Moreover, the largest percentage of cell types/states corresponded to the WNT enriched and glycolysis relating populations with the mature pancreatic ductal cell state being the minority of the populations that was further reduced with passaging (Figure S2A-B). scRNA-Seq of HPDO identified cell populations/states corresponding to pancreatic ductal cells, proliferating cells, glycolysis relating genes and processes state, a pancreatic progenitor population and a population enriched in ribosome-encoding genes (Figure 2E). Compared to PPDO, HPDO had a higher percentage of mature pancreatic ductal cell signature genes but both cultures shared the proliferating and the glycolysis signature cell states (Figure 2F&S2C). One notable difference was that the progenitor-like state of the HPDO was characterized by elevated *GP2* expression, an established marker of embryonic pancreatic progenitors^33,34^, and did not have a clear WNT signature (Figure 2F). These results correlate with the observed morphological changes, as HPDO maintain their cyst-forming phenotype, mainly characterized by mature ductal epithelial cells, while the mature ductal cell state is a minority in the PPDO that form the dense cultures upon prolonged passaging. Of note, we validated the pancreatic ductal fate of HPDO by mapping the scRNA-Seq dataset to the human tabula sapiens atlas^35^ (Figure S2D-E). Our scRNA-Seq analysis demonstrated the transcriptional similarity of different PPDO lines (neonatal compared to adult) and highlighted a potential transcriptional progenitor signature difference between PPDO and HPDO.

**Figure 2:**
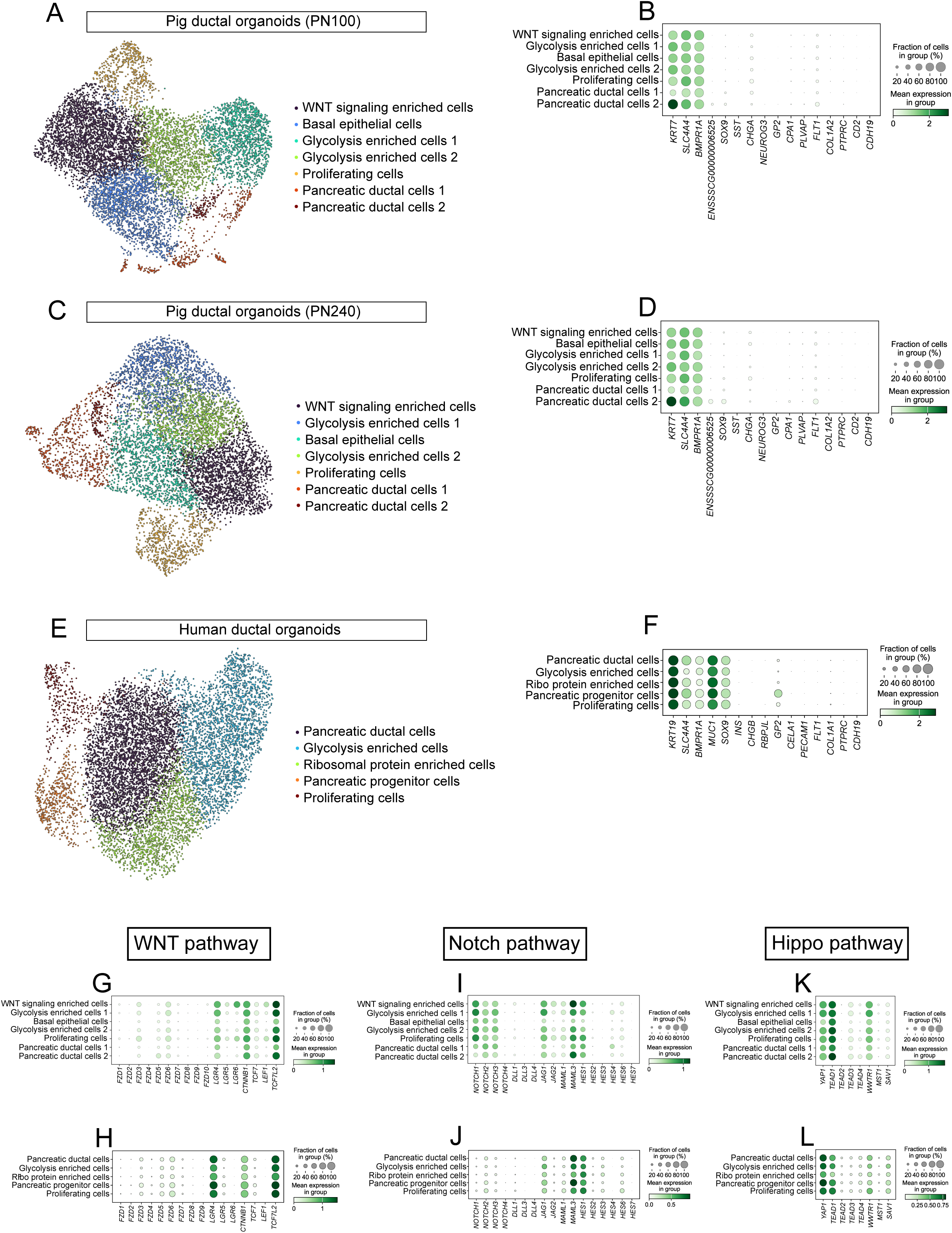
Deep-phenotyping of PPDO and benchmarking to HPDO by scRNA- Seq. (A) UMAP showing the cell types/states of scRNA-Seq data from PN100-derived PPDO at passage 7. (B) Dot plot showing expression of key pancreatic gene markers for major cell types of the PN100 PPDO scRNA-Seq dataset including ductal (*KRT7*-*SLC4A4*-*BMPR1A*- *MUC1* (*ENSSSCG00000006525*)-*SOX9*), endocrine (*SST*-*CHGA*), endocrine progenitors (*NEUROG3*), acinar (*GP2*-*CPA1*), endothelial (*PLVAP*-*FLT1*), stellate (*COL1A2*), immune (*PTPRC*-*CD2*) and schwann (*CDH19*) cell markers. (C) UMAP showing the cell types/states of scRNA-Seq data from PN240-derived PPDO at passage 4. (D) Dot plot showing expression of key pancreatic gene markers for major cell types of the PN240 PPDO scRNA-Seq dataset including ductal (*KRT7*-*SLC4A4*-*BMPR1A*- *MUC1*(*ENSSSCG00000006525*)-*SOX9*), endocrine (*SST*-*CHGA*), endocrine progenitors (*NEUROG3*), acinar (*GP2*-*CPA1*), endothelial (*PLVAP*-*FLT1*), stellate (*COL1A2*), immune (*PTPRC*-*CD2*) and schwann (*CDH19*) cell markers. (E) UMAP showing the cell types/states of scRNA-Seq data from HPDO. (F) Dot plot showing expression of key pancreatic gene markers for major cell types of the HPDO scRNA-Seq dataset including ductal (*KRT19*-*SLC4A4*-*BMPR1A*-*MUC1*- *SOX9*), endocrine (*INS*-*CHGB*), acinar/pancreatic progenitor (*GP2*), acinar (*RBPJL*- *CELA1*) endothelial (*PECAM1*-*FLT1*), stellate (*COL1A1*), immune (*PTPRC*) and schwann (*CDH19*) cell markers. (G-L) Dot plots of expression of important genes from the WNT (G-H), NOTCH (I-J) and Hippo (K-L) signaling pathways in the scRNA-Seq dataset of PN100 PPDO (G-I-K), and HPDO (H-J-L).

Using the scRNA-Seq datasets, we compared PPDO and HPDO in terms of major signaling pathways known to be active in pancreatic ductal/progenitor cells. As mentioned above, PPDO had higher expression of WNT-related genes with the prominent example of the WNT co-receptor, leucine rich repeat containing G protein-coupled receptor 6 (*LGR6*) which was enriched in the progenitor cluster compared to HPDO (Figure 2G-H). However, *LGR4* expression was uniform among cell clusters/states suggestive of distinct function for the paralogues in WNT signaling in PPDO, as all LGR protein family members bind to RSPO1^36^, which is added to the medium. Both cultures expressed key genes of the NOTCH signaling pathway, with PPDO showing an elevated expression of *NOTCH* ligands that was not the case in the HPDO cultures (Figure 2I-J). PPDO and HPDO had similar expression levels of genes involved in the Hippo pathway (Figure 2K-L). Lastly, we performed a ligand-receptor analysis to uncover intercellular signaling pathways that could be important for cellular identity and progenitor biology. For PPDO, there was a strong enrichment for epidermal growth factor (EGF) signaling ligand-receptor interactions, with additional ligand receptor pairs pointing to WNT and integrin signaling as enriched in these cultures (Figure S3A-B). The most common ligand-receptor interactions in HPDO related to integrin signaling, followed by ligand-receptor interactions in the *EGF*, (transforming growth factor (*TGF*) and mucin related pathways (Figure S3C-D). These differences are present independent of the medium composition as the same medium formulation is used for both HPDO and PPDO.

### Benchmarking PPDO to primary porcine pancreas

To benchmark PPDO to the primary tissue, we profiled porcine pancreata from two animals near reproductive maturity (PN150) using scRNA-Seq and identified all major pancreatic cell lineages (Figure S4A). First, we integrated and compared all cell types/states from the primary porcine pancreas and PPDO datasets. The two datasets did not cluster together in two-dimensional space, yet a small percentage of the PPDO cells were classified as primary pancreatic ductal cells (Figure S4B-C). Integrating only the primary ductal and beta cell populations together with all the PPDO cells showed that the primary ductal cells clustered in latent space to the pancreatic ductal cells of the PPDO, showcasing the similarities between the two populations (Figure 3A-B). To assess ductal identity, we checked the expression of the top five marker genes per PPDO cluster assigned as pancreatic ductal cells in the scRNA-Seq of the primary porcine pancreas. This comparison showed that the pancreatic ductal cell 2 population/state of PPDO was more similar to the primary ductal cells, while the pancreatic ductal cell 1 population appeared to be a culture specific cell state/type (Figure S4D-E). Importantly, we noticed that the porcine duct contains a cell population characterized by *LGR5* and anterior gradient 2, protein disulphide isomerase family (*AGR2-*recently shown to be expressed in WNT enriched population of mouse ductal cells^37^) gene expression implying a WNT responsive population with progenitor characteristics (Figure 3C-E). Immunostaining of porcine pancreas and PPDO against AGR2 indicated subpopulations of cells with AGR2 protein expression. This heterogeneity was more pronounced in the PPDO cultures (Figure 3F-G). To test the possibility that the WNT/BMP positive population gives rise to PPDO, we initiated PPDO cultures comparing complete medium and medium depleted of the WNT and BMP signaling cytokines, namely WNT3a-RSpondin1-Noggin (WRN). Initial organoid formation was unaffected in the two conditions suggesting that PPDO can form in the absence of a WNT/BMP signaling pulse (Figure 3H-I). However, the cultures without WRN factors had limited expansion potential with PPDO decreasing progressively in number and disappearing between passage three to six (Figure 3J-K). In summary, our data benchmarked the transcriptional similarities and differences of *in vitro* derived PPDO to the *in vivo* primary porcine pancreas, pointing to both WNT/BMP^+^ and WNT/BMP^-^ populations with organoid forming capacity.

**Figure 3:**
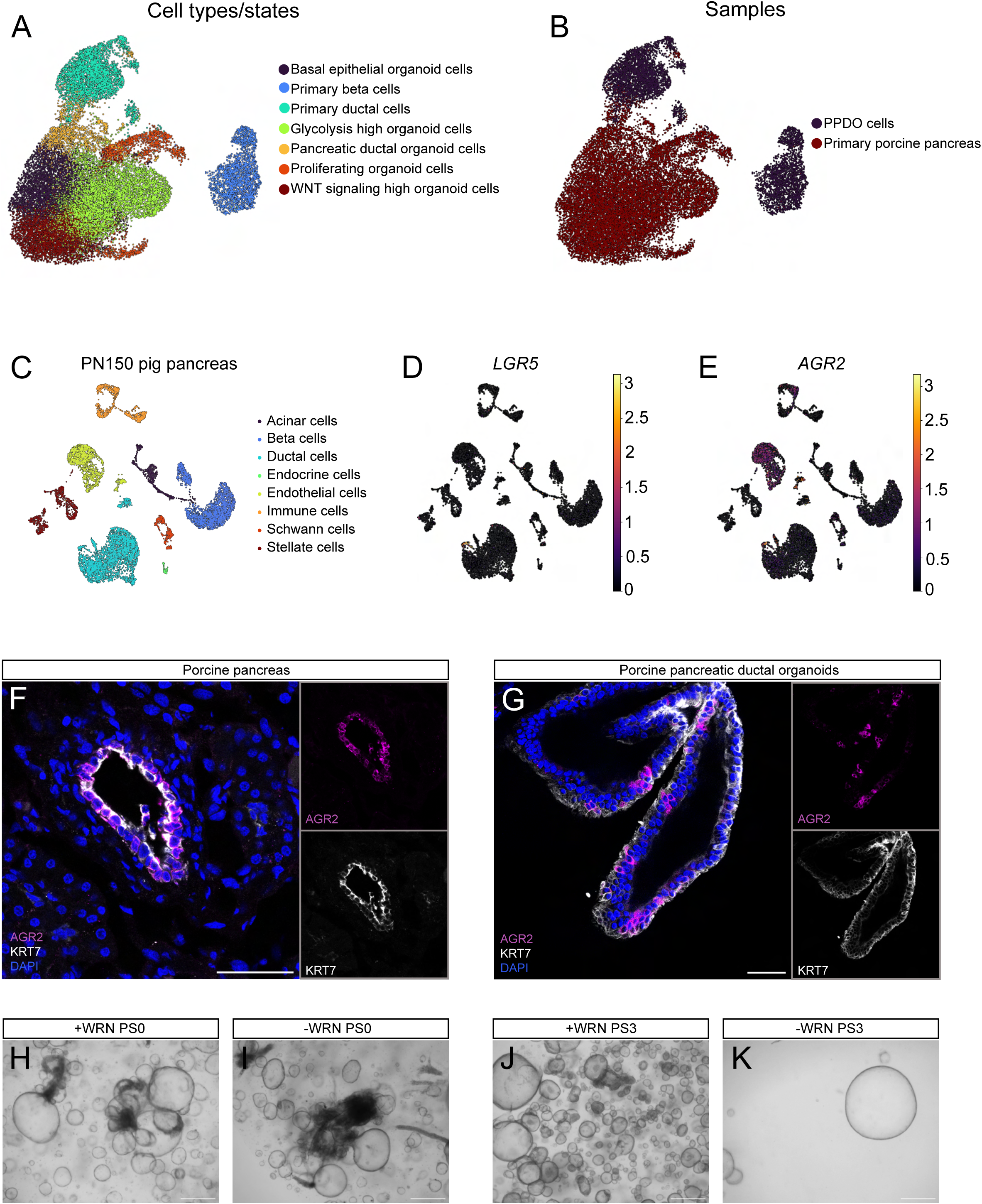
Benchmarking PPDO to primary porcine pancreas. (A-B) UMAP representation of the integrated PPDO and primary porcine pig ductal and β-cells scRNA-Seq datasets with the corresponding annotated cell types/states (A) as well as the corresponding individual samples used for integration (B). sysVI integration is shown. (C-E) UMAP representation of the PN150 pig pancreas dataset showing the corresponding cell types (E) and the expression level of the *LGR5* (F) and *AGR2* (G) genes. (F) Single-plane confocal images of porcine pancreas (PN48) immunostained against KRT7 (gray) and AGR2 (magenta) and counterstained with DAPI. Insets show the individual channels of the merge image. Scale bar: 50 µm. (G) Single-plane confocal images of PPDO immunostained against KRT7 (gray) and AGR2 (magenta) and counterstained with DAPI. Insets show the individual channels of the merge image. Scale bar: 50 µm. (H-I) Brightfield images of PPDO cultured in medium supplemented with (H) or without (I) WNT3a/R-spondin/Noggin factors (WRN) 7 days following duct isolation and culture. Scale bar 500 µm. (J-K) Brightfield images of PPDO cultured in medium supplemented with (J) or without (K) WNT3a/R-spondin/Noggin factors (WRN) after three passages at 7 days of culture post passage. Scale bar 500 µm. Experiments in (H-K) have been independently reproduced using PPDO from three different animals.

### Differentiation potential of PPDOs

Embryonic ductal epithelium contains cells with bipotent potential towards ductal and endocrine lineages and are derived from the multipotent pancreatic progenitors with tri-lineage potential (acinar-ductal-endocrine cells). To explore the utility of PPDO for studying pancreatic progenitor biology, we tested different endocrine differentiation protocols to assess PPDO’s developmental capacity towards the endocrine lineage. First, we complemented the basal organoid culture medium with porcine serum, aiming to mimic the *in vivo* milieu. Gene expression analysis did not show any significant induction of genes linked to endocrine cell fate apart from induction of neuronal differentiation 1 (*NEUROD1*) (Figure S5A-B&E-J). Next, we tested various combinations of protocols for differentiating human embryonic stem cells to β-cells, since a similar approach has been recently shown to differentiate primary mouse pancreatic organoids towards endocrine fate^29,37^. However, this approach, in particular S5 medium used for endocrine progenitor induction, was not tolerated by the PPDO, inducing massive cell death (Figure S5C-D).

Next, we tested removing the proliferating cytokines from the PPDO medium could lead to cell fate changes. To address the changes on a global scale, we performed bulk transcriptome analysis of different PPDO lines cultured for seven days in complete and cytokine-depleted media (see Methods for details). Differential gene expression analysis showed a clear transcriptional segregation of PPDO cultured in the two media. PPDO cultures from different developmental stages clustered together. This demonstrated that the developmental stage that the PPDO was derived from did not have a substantial effect on cell identity and the transcriptional signature (Figure S6A). We observed that most endocrine signature genes were not expressed in these cultures. However, we detected expression of endocrine genes in the embryonic PPDO sample, which suggested that embryonic PPDO might retain more plasticity and potency compared to adult-derived PPDO (Figure S6B). Differential gene expression analysis revealed that genes upregulated upon differentiation relate to a mature ductal gene signature, predominantly carbonic anhydrases and retinoic acid signaling pathway genes (Figure 4A). Genes in the WNT pathway were among the most significantly downregulated during differentiation, suggesting that the WNT proliferative progenitor phenotype can be dynamically regulated *in vitro* (Figure 4A). Evaluating the expression of the top 50 down- and upregulated genes in the scRNA-Seq PPDO dataset showed an enrichment of the downregulated transcripts in the WNT and glycolysis enriched clusters, while the upregulated genes were lowly expressed in this dataset, corroborating the fate transformation of PPDO (Figure S6C-D). Using pathway analysis, we identified established molecular pathways of ductal cell identity including one-carbon and retinoic acid metabolic processes among the most significantly upregulated transcripts. Pathways enriched in the downregulated genes included among others vascular endothelial growth factor (VEGF) related processes, glycolysis/hypoxia pathways and epithelial-to-mesenchymal transition processes (Figure 4B-C). Therefore, the RNA-Seq based comparison of the differentiation medium suggested that removal of proliferating agents from the PPDO medium promotes a more mature epithelial/ductal fate.

**Figure 4:**
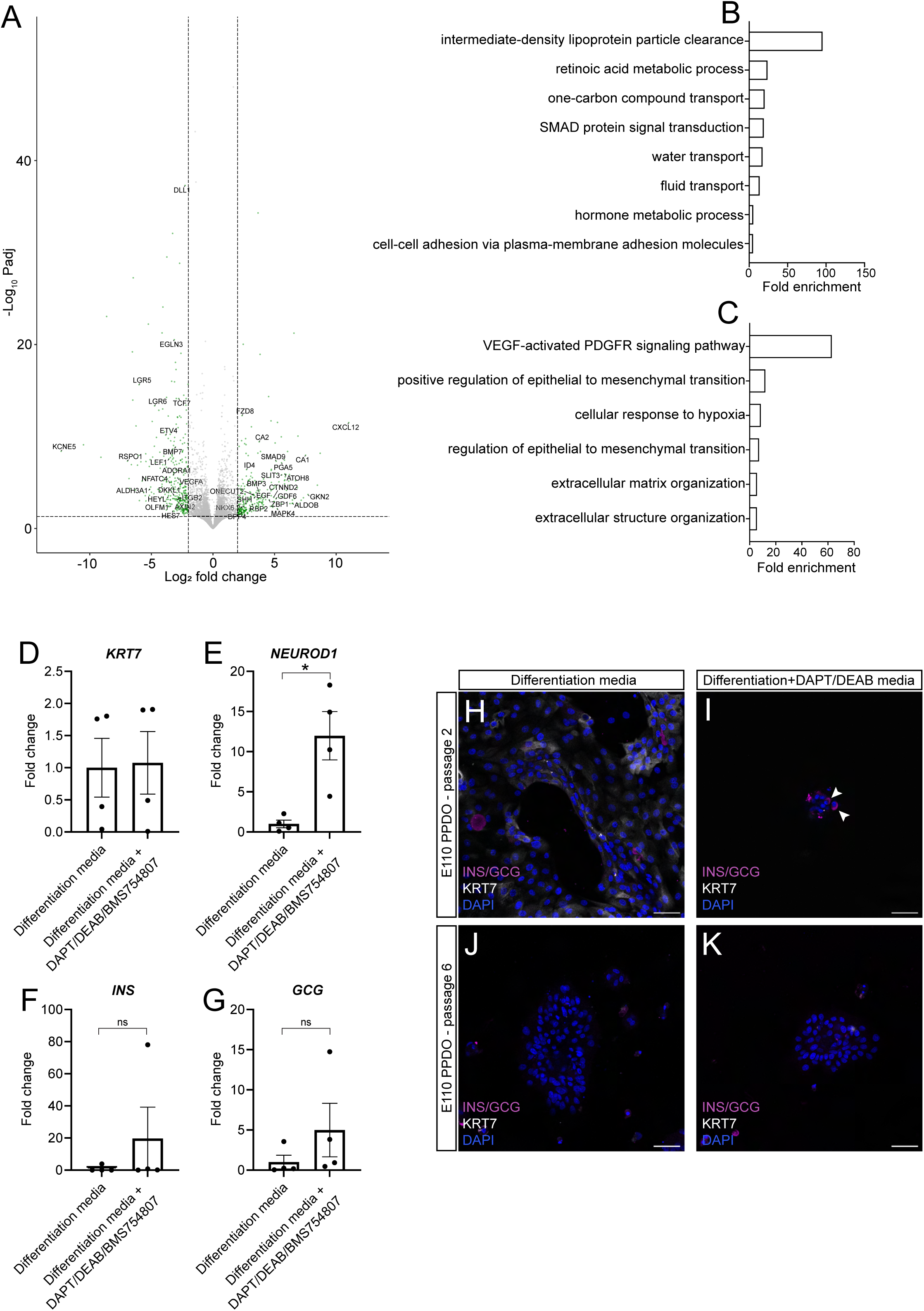
Differentiation potential of PPDO to pancreatic lineages. (A) Volcano plot indicating the number of significantly changed genes following the 7 days of culturing in the basal differentiation medium. Genes important for developmental processes are highlighted in the graph. (B-C) Bar plots showing the fold enrichment of significantly affected biological processes for the upregulated and downregulated genes of the RNA-Seq dataset shown in (A). (B) shows selected enriched GO terms for the significantly upregulated genes and (C) shows selected GO terms for the significantly downregulated genes. (D-G) Plots showing the fold change of gene expression for *KRT7* (D), *NEUROD1* (E), *INS* (F) and *GCG* (G) genes between differentiation medium alone or supplemented with DAPT-DEAB-BMS754807. Mann-Whitney test was used to assess significance with P=0.0286 for *NEUROD1*, P=0.1143 for *INS* and P=0.4857 for *GCG*. *n=4* different PPDO lines were assayed. (H-K) Single-plane confocal images of PPDO derived from E110 pig pancreas from passage 2 (H-I) or passage 6 (J-K) cultures and treated with differentiation only (H-J) or differentiation media supplemented with the NOTCH inhibitor DAPT and the aldehyde dehydrogenase inhibitor DEAB (I-K). PPDO were immunostained against KRT7 (gray)-INS/GCG (magenta) and counterstained with DAPI. Similar staining patterns were obtained from n=4 different PPDO lines. Scale bar: 50 µm.

Using the RNA-Seq results as a guide, we modified the PPDO medium to induce endocrine cell fate. We treated PPDO with a cytokine depleted media supplemented with inhibitors for NOTCH, retinoic acid and insulin receptor signaling for five days. This treatment induced gene expression of endocrine genes but with a large variability between PPDO lines (Figure 4D-G). Interestingly, PPDO derived from the late embryonic pancreas (E110) were the most amenable to endocrine induction highlighting again the differentiation potential that might be encoded in different developmental stages (outliers in Figure 4D-G). Still, no hormone expressing cells were detectable as assessed by immunocytochemistry on the protein level. To induce protein hormone expressing cells, we dispersed PPDO to single/clumps of cells (to manually disperse them from the epithelium) and plated them in laminin-coated chambers (as laminin/integrin signaling is important for endocrine differentiation during embryogenesis^38–40^), followed by treatment with the endocrine induction medium. Of note, the insulin signaling inhibitor was omitted as it was inducing overt cell death over prolonged treatment. Under these conditions, we observed few scattered INS^+^/GCG^+^ (co-stained for both markers) cells in early but never in late passaged PPDO (Figure 4H-K). Our results showcase the utility of PPDO as a platform to study ductal cell maturation *in vitro* and highlight a potential epigenetically-encoded variability towards induction to endocrine fate.

### Chemical screen identifies regulators of PPDO proliferation/survival

Organoid cultures hold great translational potential for drug discovery/repurposing studies. We designed and performed a PPDO-based chemical screen to reveal FDA-approved drugs that could be a potential safety risk for ductal/progenitor cells by inducing proliferation (carcinogenesis) or cell death (pancreatitis). We plated PPDO in 96-well plates, cultured for two days for PPDO formation and treated them for three days with 176 chemicals from a library of FDA-approved (see methods for screening details). We used high-content imaging to identify chemicals that induced toxicity or mitosis as assayed by positivity for the mitosis marker phosphorylated serine 10 of histone 3 (pHH3) (Figure 5A). Brightfield imaging showed a total of 31 compounds that almost completely killed PPDO (Figure 5B-E). Grouping the toxic chemicals according to their proposed mechanism of action, we observed a clear enrichment of chemicals that inhibit DNA synthesis relating processes possibly by inhibiting cell proliferation (Figure 5F). Other prominent chemical classes inducing toxicity included adrenergic receptor inhibitors, 3-hydroxy-3-methyl-glutaryl-CoA reductase (HMG-CoA) inhibitors (statins) and estrogen receptor targeting molecules (Figure 5F). These results revealed FDA-approved chemicals with toxicity to porcine pancreatic duct/progenitor population.

**Figure 5:**
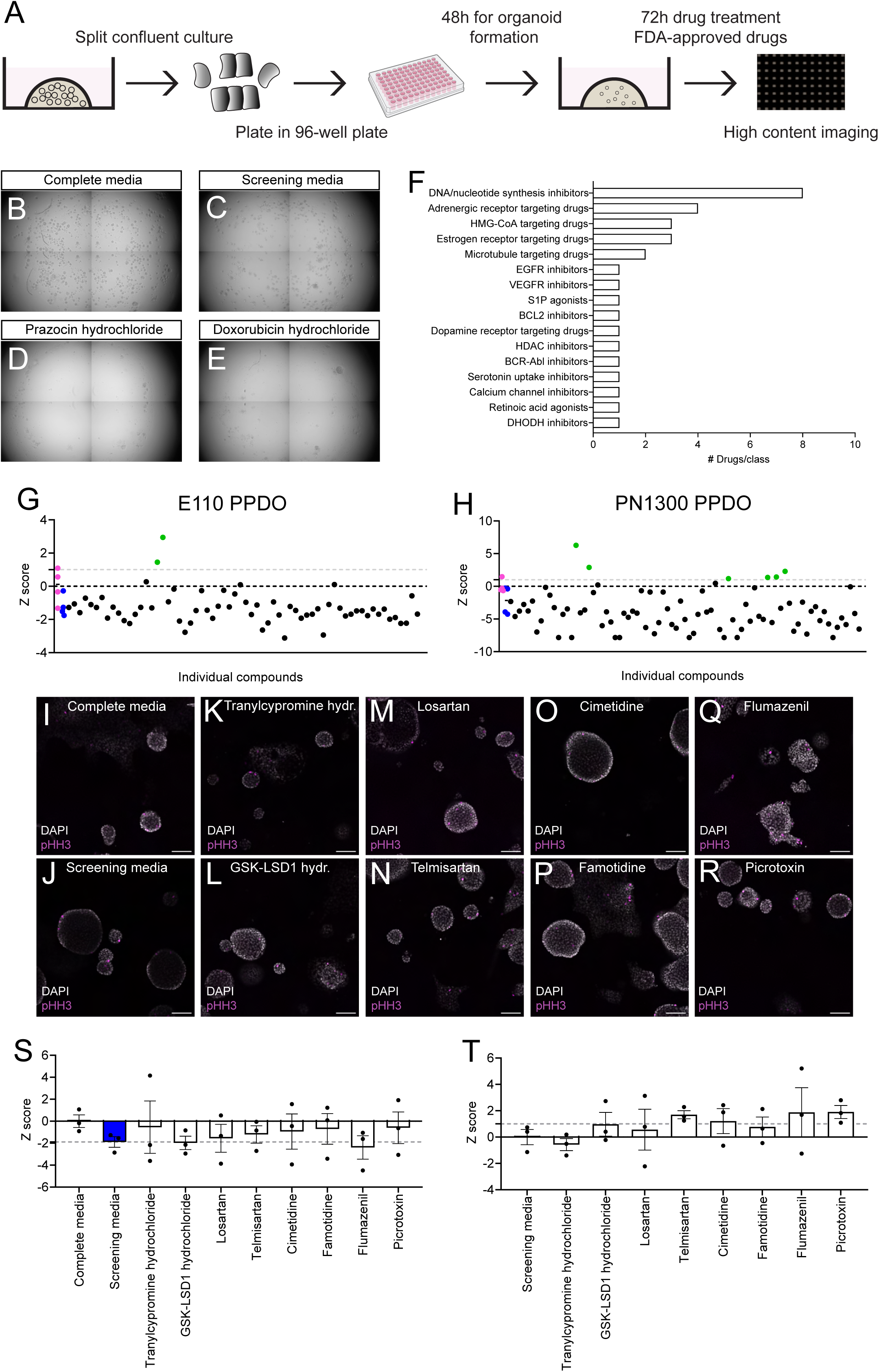
Chemical screen identifies regulators of PPDO proliferation/survival. (A) Schema documenting the chemical screen approach using PPDO cultures. (B-E) Brightfield images of the whole well of a 96-well plate picturing PPDO in complete medium (B), screening medium (C) and PPDOs treated with prazosin hydrochloride (D) and doxorubicin hydrochloride (E), two drugs that induced cell death. Four individual images were stitched together to get the overview of the well. (F) Bar plot showing the summary of the number of chemicals/chemical class that induced toxicity in the primary screen. (G-H) Scatter plots reporting the z-scores of the proliferation percentage of the primary screen results compared to the complete medium control wells for two PPDO lines, E110 (G) and PN1300 (H). Each dot represents a single well of the 96-well plate treated with a single chemical. Magenta dots show the individual wells of PPDOs treated in the complete medium with DMSO (control) and blue dots show the individual wells of PPDOs treated in screening medium with DMSO and the line for these two wells corresponding to the median of the four wells/condition. Gray dotted line demarcates z-score of 1. (I-R) Single-plane confocal images of PPDOs from the secondary screen cultured in complete (I) or screening media (J-R) and treated with DMSO (I-J), tranylcypromine hydrochloride (K), GSK-LSD1 hydrochloride (L), losartan (M), telmisartan (N), cimetidine (O), famotidine (P), flumazenil (Q) and picrotoxin (R). Proliferating cells are marked by the marker pHH3 (magenta) and nuclei are counterstained with DAPI (gray). Scale bar 100 µm. (S-T) Bar plots reporting the z-scores of the proliferation percentage of the secondary screen. In (S) the z-score of the treatments of the chemicals added to the complete medium in reference to the complete medium are shown. In (T) the z-score of the treatments of the chemicals added to the screening medium in reference to the screening medium are shown. Gray dotted line shows the mean of the screening medium z-score for (S) and the z-score of 1 for (T). *n*=3 individual PPDO lines. Error bars show the ±SEM.

On the opposite spectrum, we aimed to discover chemicals that could stimulate proliferation of PPDO cells i.e have a Z-score of zero or larger. Our primary screen showed that indeed removal of WRN factors from the medium reduced proliferation (Figure 5G-H). Eight chemicals exhibited a z-score >1 suggesting that they might stimulate proliferation compared to the complete medium (Figure 5G-H). Primary hits of our chemical screen were: flumazenil (γ-aminobutyric acid type a receptor (GABAa) receptor antagonist), cimetidine (histamine receptor H2 (HRH2) antagonist), tranylcypromine hydrochloride (lysine demethylase 1A (LSD1) inhibitor), megestrol acetate (progesterone receptor agonist), stavudine (nucleoside analog) and losartan/telmisartan/candesartan (angiotensin AT1 receptor (AGTR1) antagonist). To validate the results of the primary screen, we performed a secondary screen with the most promising primary hits using three independent PPDO lines derived from different pig developmental ages. Additionally, we included a second chemical targeting the same protein to validate the molecular pathway of the potential hits (Figure 5I-R). Our secondary screen revealed that none of the primary hits could significantly increase proliferation of PPDO when added to complete medium (Figure 5S). The GABAa receptor antagonists, flumazenil and picrotoxin could revert proliferation levels almost to the level of the complete medium, but with great variability between PPDO lines, and therefore the results did not reach statistical significance (Figure 5T). Last, we validated that the molecular targets of the primary hits were indeed expressed in the PPDO (Figure S7A-B). Overall, our chemical screen using PPDO identified drugs with potential safety issues for pancreatic ductal/progenitor cells in WRN depleted media.

### HMG-CoA and alpha-adrenergic receptor inhibitors induce toxicity in HPDO

We tested the most consistent drug phenotypes from our PPDO chemical screen for their translation potential to the HPDO system. First, we validated the gene expression of the chemical targets from the increased proliferation phenotype in our HPDO scRNA-Seq dataset and confirmed expression for the genes in the histamine receptor, GABA receptor and LSD1 demethylase targets but not for the angiotensin receptor genes, and therefore the respective chemicals were excluded from downstream experiments (Figure 6A). We screened three HPDO lines with the same chemicals inducing mitosis in the PPDO as well as with three randomly chosen chemicals from the top chemical categories that induced toxicity of PPDO cultures (Figure 5F). The estrogen receptor modulator, raloxifene, did not exhibit the same toxicity level as in the PPDO but lovastatin treatment induced a high-degree of cell death in the HPDO cultures (Figure 6B-E). Prazosin (α-adrenergic receptor blocker) was the most potent inducer of cell death HPDO cultures, a phenotype that was not prominent with carvedilol treatment, which has higher affinity for the β-adrenergic receptor (Figure 6F-G). Compared to the PPDO, HPDO showed great variability in the incorporation of the mitotic marker pHH3 across lines highlighting an inter-donor variability of proliferation dynamics. In terms of HPDO proliferation, we observed a strong influence of WRN factors as removal of these cytokines drastically diminished the percentage of pHH3^+^ cells (Figure 6H-I&R). None of the chemicals altered the proliferation percentage when added to the complete medium, with a trend for lovastatin and prazosin treatments to reduce proliferation (Figure 6R). None of the chemicals reverted the proliferation rate of HPDO to the control condition, unlike what we observed with the GABAa receptor antagonists in the PPDO cultures. Yet, picrotoxin almost tripled the proliferation rate compared to the screening medium (Figure 6H-Q&S). Our data underlined the translational potential of the PPDO for pancreas biology and pinpoint to a potentially conserved role for HMG-CoA and α- adrenergic receptor inhibitors in mammalian pancreatic ductal/progenitor cell toxicity.

**Figure 6:**
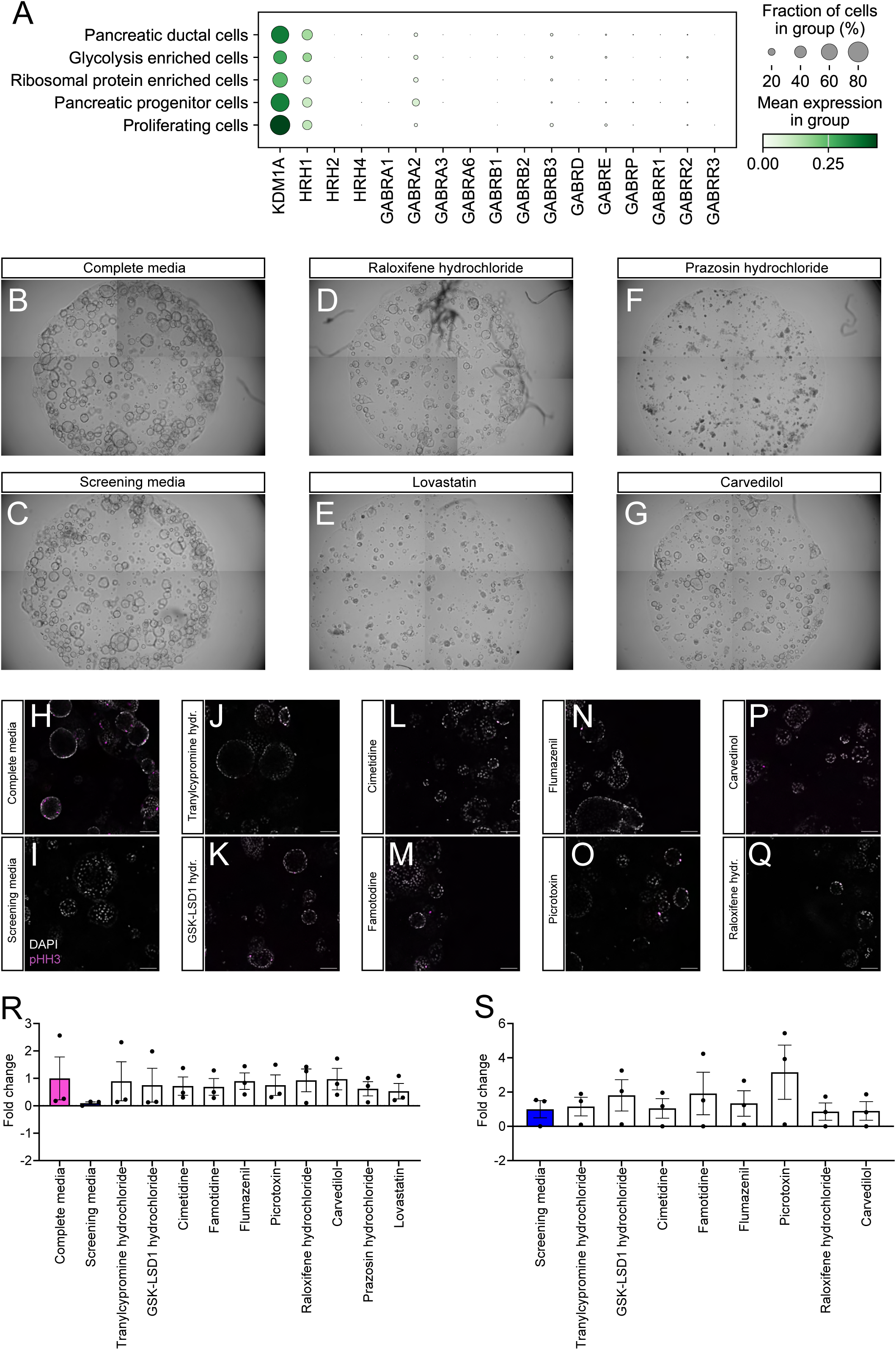
HMG-CoA and alpha-adrenergic receptor inhibitors induce toxicity in HPDO. (A) Dot plot showing gene expression of genes encoding for proteins targeted by the positive hits in the HPDOs scRNA-Seq dataset. (B-G) Brightfield images of the whole well of a 96-well plate picturing organoids in complete medium (B), screening medium (C) and PPDOs treated with raloxifene hydrochloride (D), lovastatin (E), prazosin hydrochloride (F) and carvedilol (G). Four individual images were stitched together to get the overview of the well. (H-Q) Single-plane confocal images of HPDOs cultured in complete (H) or screening medium (I-Q) and treated with DMSO (H-Q), tranylcypromine hydrochloride (J), GSK- LSD1 hydrochloride (K), cimetidine (L), famotidine (M), flumazenil (N), picrotoxin (O), raloxifene hydrochloride (P) and carvedilol (Q). Proliferating cells are marked by the pHH3 (magenta) and nuclei are counterstained with DAPI (gray). Scale bar 100 µm. (R-S) Bar plots reporting the fold change of the proliferation percentage of the HPDO chemical screen. In (R) fold change in reference to the proliferation rate of the complete medium is shown. In (S) fold change in reference to the proliferation rate of the screening medium is shown. *n*=3 individual HPDO lines. Error bars show the ±SEM.

## Discussion

Organoid cultures are a valuable tool for studying adult progenitor biology and pancreatic disease. In this work, we established and performed an in-depth characterization of PPDO across distinct developmental stages of porcine pancreas development. These samples provide information on the full range of pancreas development in a relevant large animal model (pig), samples that are difficult to obtain from human donors. Our phenotyping reveals a WNT enriched signaling progenitor signature *in vitro* that correlates with a porcine ductal subpopulation *in vivo*. No morphological or transcriptional differences were observed in PPDO from distinct developmental stages, i.e. fetal, early post-natal and adult pancreas, except from a slight improved endocrine differentiation potential of embryonic PPDO. Contrary to HPDO that maintained a similar phenotype across cultures, PPDO transformed to dense, WNT signaling enriched spheres upon prolonged culturing highlighting a difference between the two species. Moreover, we show that PPDO can acquire either a more mature ductal fate or differentiate/transdifferentiate to an endocrine fate upon chemical and extracellular matrix manipulations. Finally, we demonstrate the translation potential of this system for FDA-approved drug safety application by identifying novel conserved pathways for ductal/progenitor cell toxicity/proliferation between pigs and humans.

PPDO transcriptional and functional phenotyping confirms the ductal lineage of these cultures that was progressively lost over passaging in favor of a low proliferating, WNT signaling enriched population/state. Two studies recently reported the derivation of PPDO for the study of exocrine-related pathologies corroborating the initial pancreatic ductal fate of these cultures^41,42^. Contrary to PPDO, HPDO cultures retained their ductal fate over passaging as evident by our scRNA-Seq datasets in line with a recent scRNA-Seq study of HPDO^43^. This species difference could be explained by diverse starting culture populations or variable growth factor efficacies in the PPDO cultures. Additionally, PPDO originating from distinct porcine developmental stages do not vary morphologically, functionally, or transcriptionally, which differs to mouse intestinal organoids that show a clear morphological difference between embryonic and adult cultures^44^. Importantly, removal of proliferation-related cytokines from PPDO induces a mature ductal fate, but all endocrine-inducing protocols tested were largely inefficient. This limited endocrine potential of PPDO differs from the endocrine induction capacity of early postnatal porcine islet cultures, in which ductal cells differentiate to endocrine cells with high efficiency^13,45–48^. However, we highlight a tendency for embryonic PPDO to have increased plasticity when challenged with differentiation protocol, a plasticity that could be epigenetically encoded. Overall, we have performed a deep-phenotyping of PPDO according to recent suggestions for benchmarking organoid cultures^16^ and we argue that a cross-species comparisons in terms of molecular profile and differentiation capacity/conditions is needed to provide a holistic overview of these models^49^.

We generated PPDO across developmental stages with the aim to temporally resolve and study adult pancreatic progenitor biology, a much-debated topic in pancreas regeneration field^50^. Since expansion of pancreatic progenitors is a much-sought advancement that the field is working for mass endocrine cell production^23,51,52^, PPDO can serve as a model to study progenitor dynamics. Our results indicate a WNT enriched transcriptional signature present in both PPDO and primary porcine ductal cells. Two studies proposed that a WNT responsive population characterized by LGR5 expression could be surfacing under injury/high metabolic demand conditions^17,53^. A recent scRNA-Seq study of the adult mouse ductal epithelium clearly showed a WNT responsive transcriptional signature in the healthy mouse pancreas^37^, a population that, if present, was not highlighted in another mouse ductal scRNA-seq dataset^54^. No definitive WNT enriched population was shown/discussed in recent transcriptional profiles of human embryonic pancreas^55,56^ or in a scRNA-Seq of adult human ductal cells^10^ but that could be attributed to the enrichment strategy chosen in the latter study. A recent finding suggests that LGR5^+^ marks a multipotent progenitor population in the human embryonic pancreas, but the prevalence of this population in the scRNA-Seq datasets of human embryonic progenitors is not clear^57^. Nevertheless, these human embryonic organoids differentiated to all pancreatic epithelial lineages, similar to observations from mouse embryonic explant cultures^57,58^. Moreover, our scRNA-Seq of HPDO did not show such pronounced WNT transcriptional signature but rather progenitors marked by *GP2* expression, a multipotent embryonic progenitor marker in humans, highlighting another important difference to the PPDO^33,34,59^. Still, this population expressed hallmarks of WNT responsive cells including olfactomedin 4 (*OLFM4*) and *LGR5*^60,61^. Our data hint towards a WNT signaling dependent adult pancreatic progenitor population with organoid forming capacities, unipotent ductal progenitor potential and limited endocrine differentiation potential.

Chemical screens using organoids as a platform hold great promise for identifying new therapeutic leads or assessing the safety profile of already commercialized drugs. We utilized our PPDO system and performed a chemical, high-content imaging-based screen to assess the safety of FDA-approved chemicals for pancreatic ductal/progenitor cells. Given the need for non-rodent animal model research for regulatory approval of drugs, our PPDO platform can be utilized to screen for pancreatitis or pancreatic cancer drug applications. Our approach implicates adrenergic receptor inhibitors and HMG-CoA inhibitors in ductal/progenitor cell toxicity. While neurotransmitters are implicated in pancreatic cancer biology, no extensive studies have been performed using the α adrenergic receptor inhibitor^62^. Angiotensin-converting enzyme inhibitors (ACEi) have been involved in drug-induced pancreatitis previously, suggesting that this pathway can be involved in pancreas damage^63,64^. On the contrary, there is a wealth of clinical data regarding potential roles of statins in exocrine disease such as pancreatitis and pancreatic cancer^65–67^, and our study points to ductal cells as a target for this drug class. Whether statins can induce toxicity in patient-derived pancreatic ductal adenocarcinoma organoids remains to be seen. Moreover, our screening approach highlights the absence of significant proliferative phenotypes for FDA-approved drugs, but a minor pro-proliferative effect of GABAa receptor inhibitors. Of note, GABA treatment itself has been previously involved in pancreatic β-cell proliferation and in contradictory studies regarding α- to β-cell fate conversion^68–74^. The absence of proliferative phenotypes is of importance for pancreatic cancer safety concerns in the presence of pre-cancerous lesions and highlights the use of the easily obtainable PPDO for this type of studies.

In summary, we performed an in-depth benchmarking of PPDOs across developmental stages and used them as a platform for high-content chemical screen. The wider accessibility of pig pancreas tissue across development compared to human and the validation of PPDO as a screening platform with comparable results to HPDO makes it an attractive model for the study of mammalian ductal/progenitor biology at scale and for pancreatitis/pancreatic cancer applications. Meanwhile, we acknowledge that biological results using pancreatic organoids can vary greatly depending on the i) starting culture material (primary vs stem cell derived), ii) developmental stage (embryo vs adult), iii) model organism (mouse vs pig vs human) and iv) culture conditions (medium composition/passaging). Thus, clear reporting of such parameters will be necessary to clarify the biologically relevant results. Better characterized pancreatic organoid systems can pave the way for a holistic understanding of pancreatic biology and disease, and generation of new therapy approaches.

## Resource availability

Raw and processed sequencing data will be available upon publication of the research article. Code used to analyze scRNA-Seq and bulk RNA-Seq data are available and will be updated here: https://github.com/chrika2/Pig_organoids. All data in the manuscript are available upon request from the corresponding author.

## Acknowledgements

We would like to thank Kerstin Diemer, Susanne Badeke, Ines Kunze and Christina Blechinger for their excellent technical support as well as Drs Alessandro Dema, Mostafa Bakhti and Elke Schlüssel for critical comments on the manuscript. We acknowledge the technical support of the Genomics Core Facility at Helmholtz Munich. Next-Generation-Sequencing was also carried out at the Competence Centre for Genomic Analysis (Kiel). C.K. was supported by a postdoctoral fellowship from the Alexander von Humboldt foundation. M.M.v.D.B was supported by the Helmholtz Research School for Diabetes (HRD), which is funded by the Helmholtz Association - Initiative and Networking Fund (IVF). The study was supported by the Helmholtz Society, Deutsche Forschungsgemeinschaft (DFG, German Research Foundation) – Project number 458958943, DFG Research Infrastructure NGS_CC (project 407495230) as part of the Next Generation Sequencing Competence Network (project 423957469), the German Center for Diabetes Research (DZD; 82DZD08D03) and the Juvenile Diabetes Research Foundation (Breakthrough T1D; 3-SRA-2023-1420-S-B).

## Author contributions

C.K conceptualized the study, performed and analyzed most of the experiments and wrote the first draft of the manuscript. K.Y generated and characterized embryonic porcine organoid cultures, assisted in computational analysis and provided intellectual input on study design. M.S prepared scRNA-Seq samples and assisted in computational analysis. M.M.V.d.B assisted with porcine pancreas preparations. T.K prepared and provided human tissue. J.F and S.F processed assisted in scRNA-Seq project design and processed NGS data. E.K, S.R and E.W provided intellectual input, prepared porcine pancreas samples and were responsible for pig colony maintenance. H.L contributed to study design, provided financial support and supervised the entire study. All authors reviewed and edited the manuscript.

## Supplementary Figure legends

**Supplementary Figure 1:**
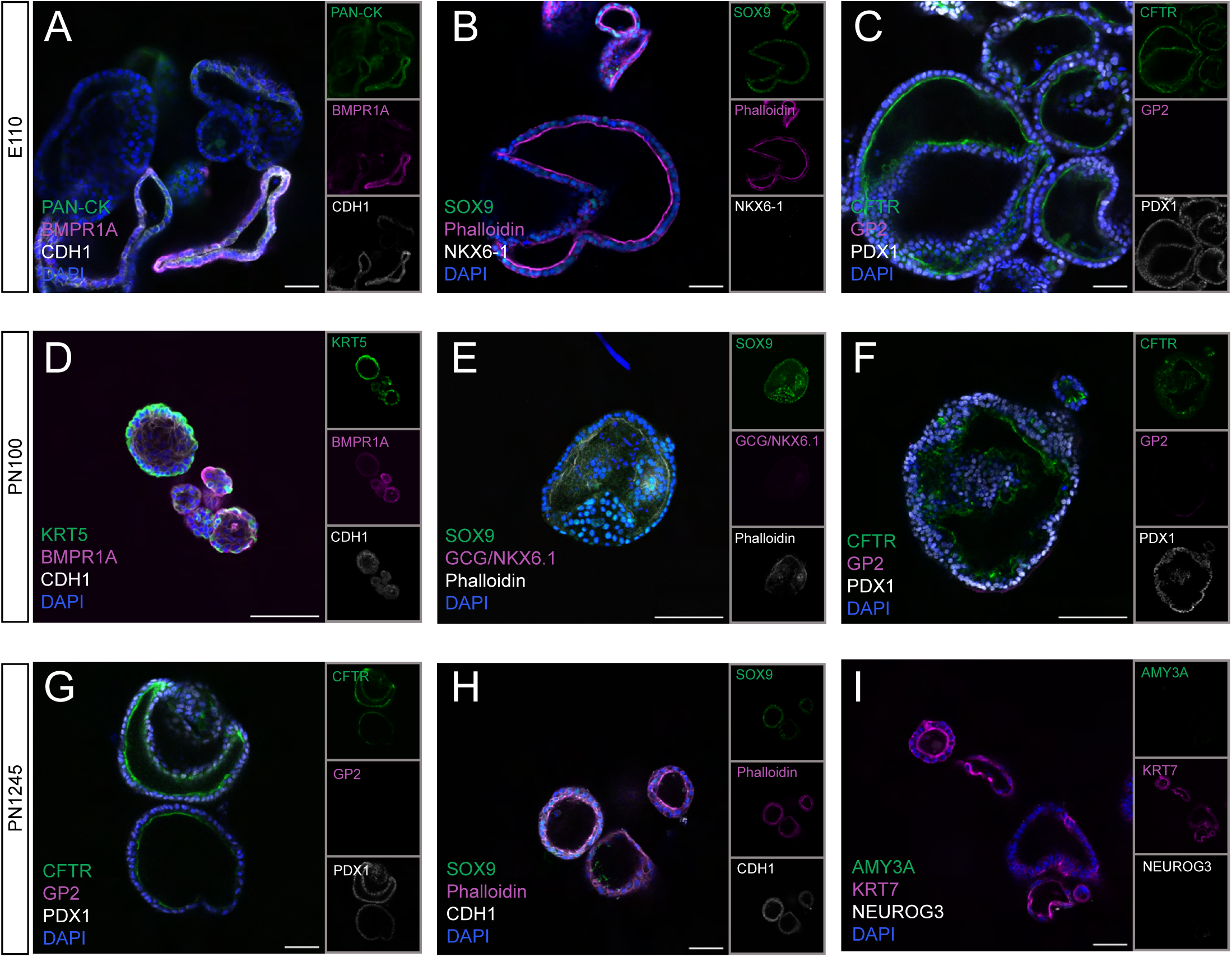
Pancreatic marker expression in PPDO from different developmental stages. (A-C) Single-plane confocal images of PPDO derived from E110 pig pancreas immunostained against Pan-Cytokeratin (PAN-CK)-BMPR1A-CDH1 (A), SOX9- Phalloidin-NKX6-1 (B), CFTR-GP2-PDX1 (C) and counterstained with DAPI. Insets show the individual channels of the merge image. Scale bar: 50 µm. (D-F) Single-plane confocal images of PPDO derived from PN100 pig pancreas immunostained against KRT5-BMPR1A-CDH1 (D), SOX9-GCG/NKX6-1-Phalloidin (E), CFTR-GP2-PDX1 (F) and counterstained with DAPI. Insets show the individual channels of the merge image. Scale bar: 50 µm. (G-I) Single-plane confocal images of PPDO derived from PN1245 pig pancreas immunostained against CFTR-GP2-PDX1 (G), SOX9-Phalloidin-CDH1 (H), AMY3A- KRT7-NEUROG3 (I), and counterstained with DAPI. Insets show the individual channels of the merge image. Scale bar: 50 µm.

**Supplementary Figure 2:**
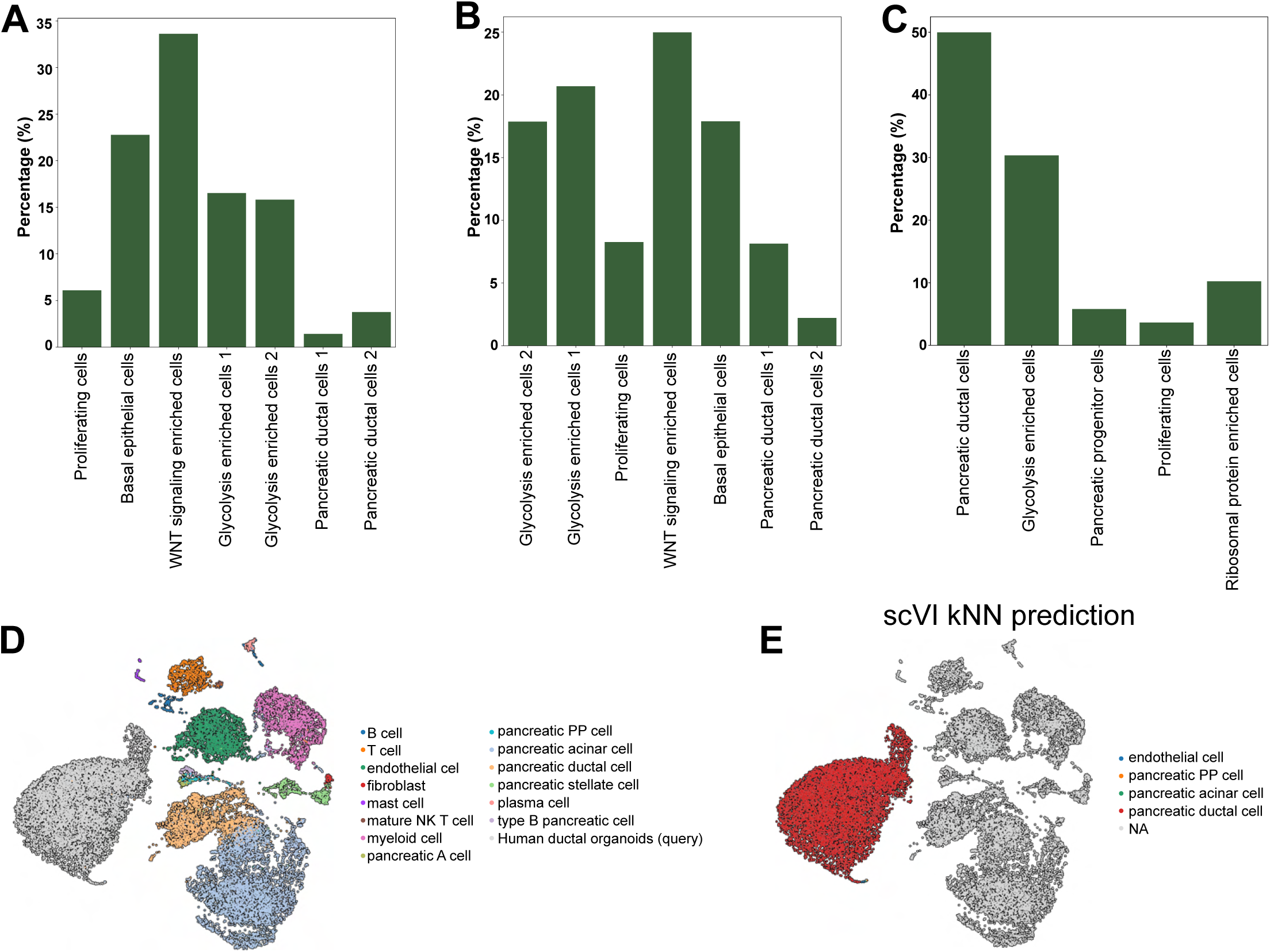
Cell type/states classification of PPDO/HPDO. (A-C) Bar-plots showing the percentage of each cell state/type of the scRNA-Seq in PN100 (A) and PN240 (B) PPDO as well as in HPDO (C). (D-E) UMAP representation of the cell label transfer of the HPDO form the human tabula sapiens pancreas dataset. (D) depicts all the cell types of the atlas together with the query cells and (E) is the predicted cell labels to the query dataset using the scVI model.

**Supplementary Figure 3:**
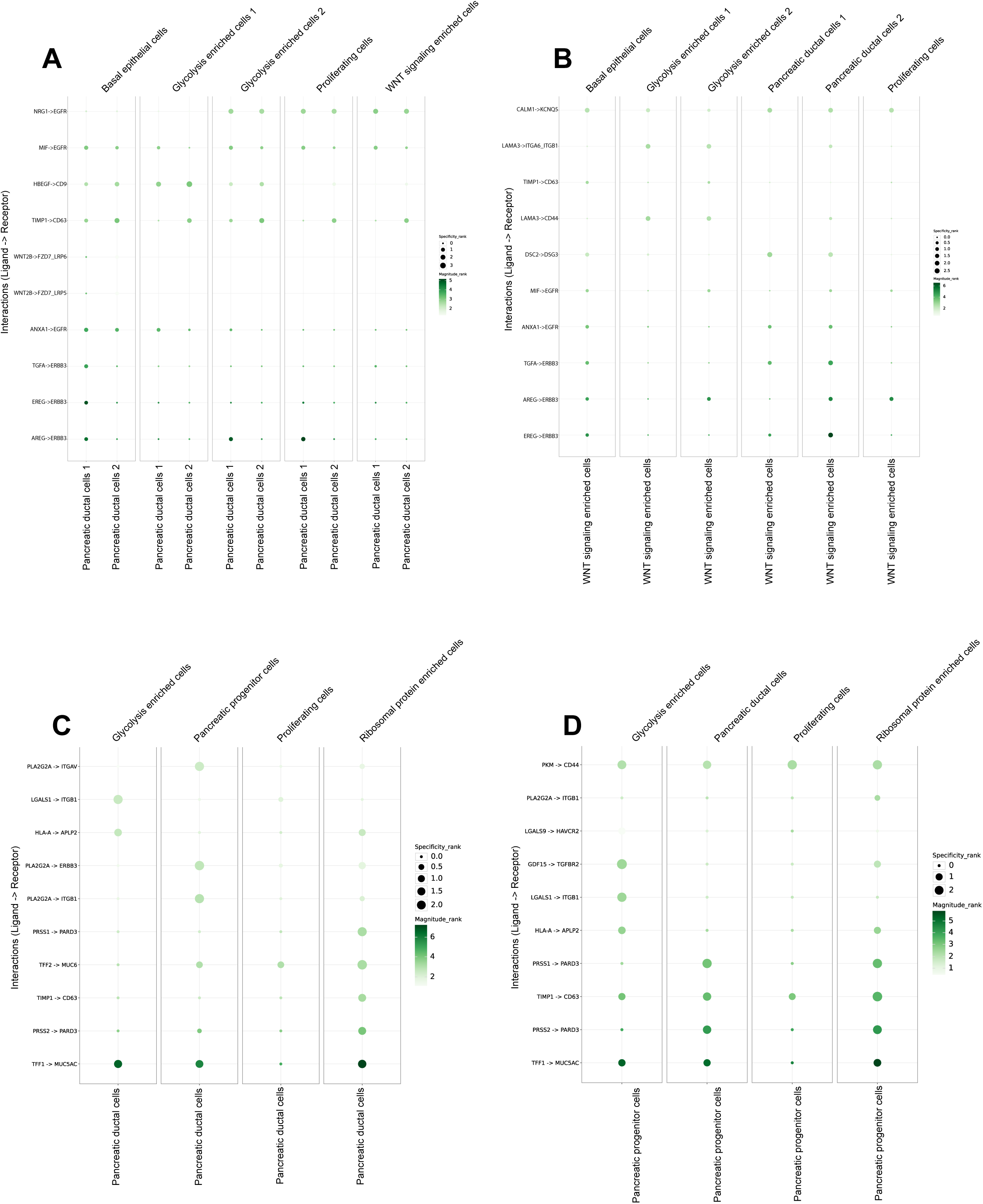
Ligand-receptor interactions in PPDO/HPDO. (A-D) Dot plots showing the most significant enriched ligand-receptor interactions towards ductal (A) and progenitor cells (B) in PPDO and (C-D) HPDO scRNA-Seq data respectively.

**Supplementary Figure 4:**
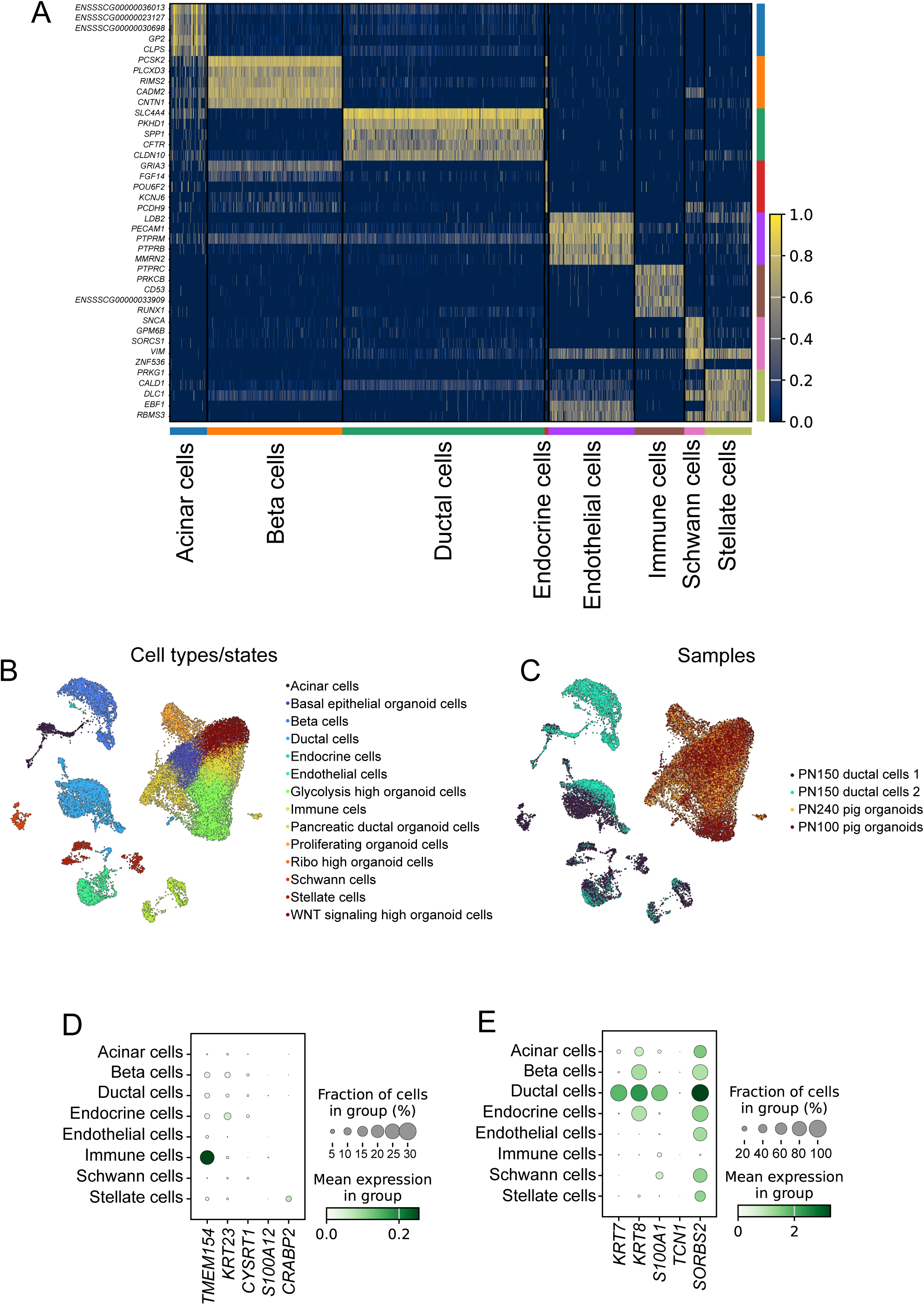
Characterization of PN150 primary porcine dataset. (A) Heatmap showing the top 5 marker genes per cell type identified in the scRNA-Seq data of 2 PN150 porcine pancreata. (B-C) UMAP representation of the integrated PPDO and primary porcine pig pancreas scRNA-Seq datasets with the corresponding annotated cell types/states (B) as well as the corresponding individual samples used for integration (C). Scanorama integration is shown. (D-E) Dot plot showing the expression of the top 5 gene markers from the pancreatic ductal cell 2 (D) and pancreatic ductal cell 1 (E) PPDO cell types/states in the primary porcine pancreatic dataset.

**Supplementary Figure 5:**
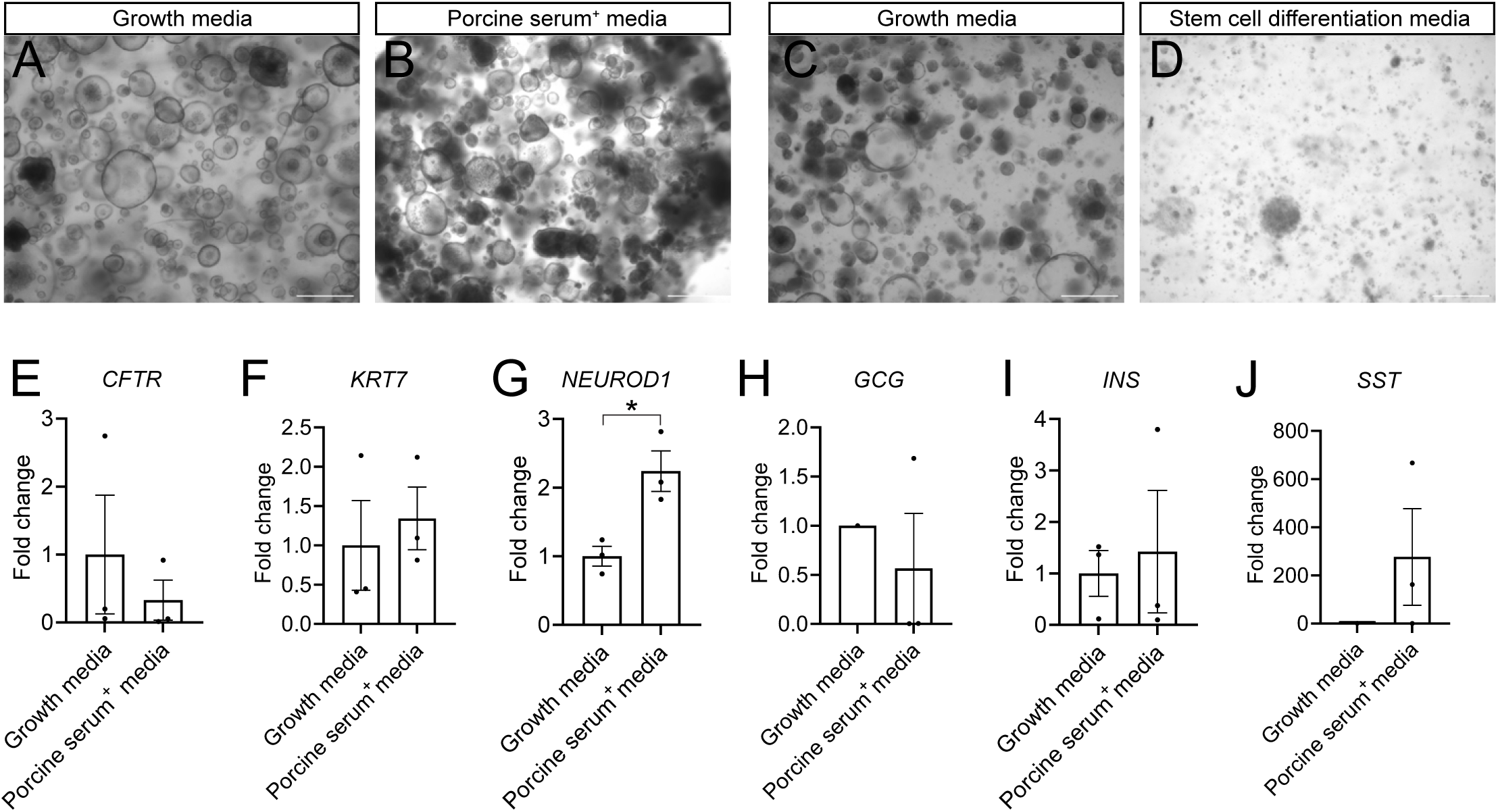
Endocrine differentiation protocol testing. (A-B) Brightfield microscopy images of PPDO in complete (A) or porcine serum supplemented (B). Scale bar 500 µm. (C-D) Brightfield microscopy images of PPDO at the end of differentiation in complete (C) or at the end of the differentiation using S5+S6 combination (D) media. Scale bar 500 µm. (E-J) Bar plots showing the fold change of gene expression analysis at the end of the differentiation after treatment with porcine serum. Gene expression was measured for *CFTR* (E), *KRT7* (F), *NEUROD1* (G), *GCG* (H), *INS* (I) and *SST* (J). n=3 independent PPDO lines. Absence of samples from the plots indicate non-detectable amplification following the RT-PCR.

**Supplementary Figure 6:**
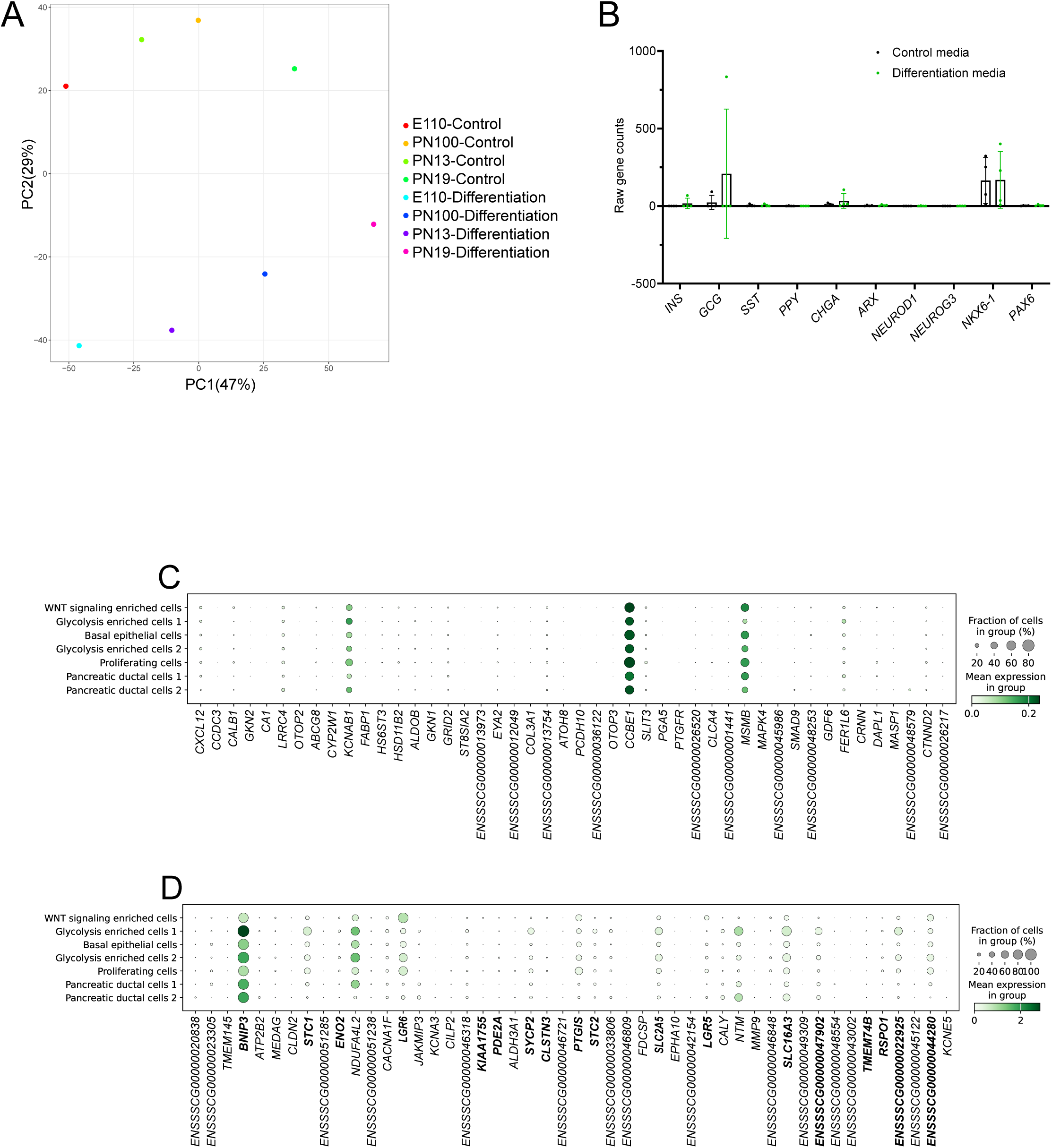
Quality assessment of the RNA-Seq differentiation analysis. (A) PCA plot of the bulk RNA-Seq results from the differentiation test showing the first two components. (B) Bar plot showing the raw counts (gene length scaled) of the RNA-Seq dataset for control and differentiation media. Each dot represents a different PPDO line sequenced (n=4). (C-D) Dot plots showing the expression of the initial top 50 significantly upregulated (C) and downregulated genes (D) in the scRNA-Seq dataset of the PN100 PPDO sample. Some genes were not expressed in the scRNA-Seq dataset and therefore are not present in the dot plot.

**Supplementary Figure 7:**
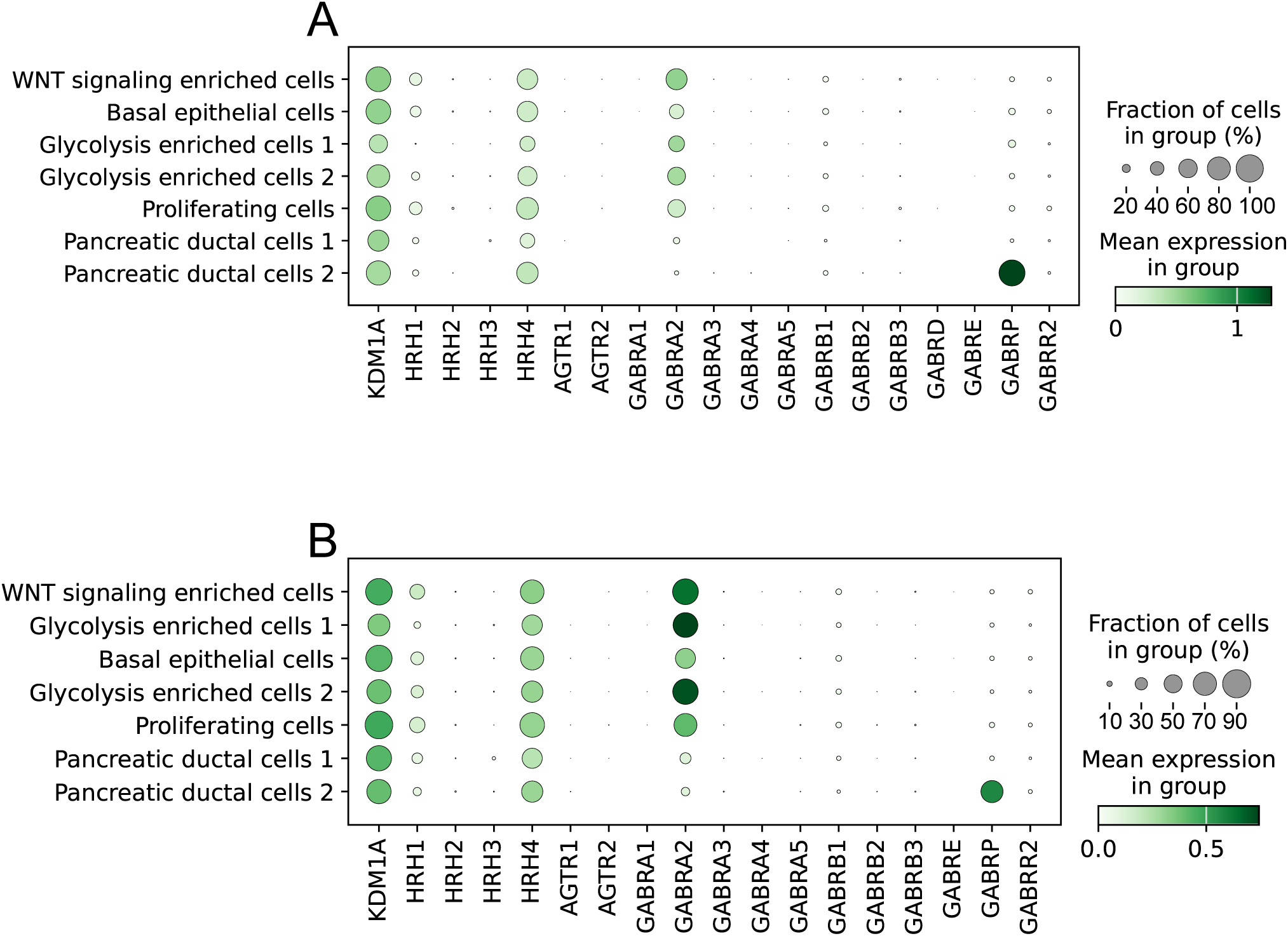
Expression of chemical screen targets in PPDO. (A-B) Dot plots showing gene expression of the genes in the pathways targeted by the primary hits of the chemical screen in the PN100 (A) and PN240 (B) scRNA-Seq dataset.

## Materials and methods

### Porcine pancreatic ductal organoid culture

Wild-type, *INS*-eGFP and *INS*-eGFP;*INS*:C94Y Landrace pigs were housed at the state-of-the-art, pathogen-free pig facility of LMU Munich (Center for Innovative Medical Models; www.lmu.de/cimm/). Work described in this paper was performed under the LMU, Permission No. 55.2-2532.Vet_02-17-136 and 55.2-2532.Vet_02-19-195. All experiments were conducted according to the German Animal Welfare Act and Directive 2010/63/EU on the protection of animals used for scientific purposes.

To establish PPDO, the isolated pig pancreas was dissected, cut into small pieces and digested with 1mg/ml collagenase V (diluted with HBSS 1X buffer) for 20’ at 37°C. Digestion reaction was stopped with 1% BSA containing buffer in PBS 1X and ductal structures were manually picked under the microscope. Ductal structures were washed five times with what we refer to as basal medium constituted of Advanced DMEM/F12 (Life Technologies) medium supplemented with Pen/Strep (1X), primocin (100µg/ml), GlutaMAX (1X) and HEPES (100mM). PPDO were maintained with the previously published complete medium for HPDO^20^ containing the basal medium supplemented with: N2 supplement (1X – ThermoFischer Scientific), B27-vitamin A supplement (1X – Life Technologies), A83-01 (5µM – Tocris Biosciences), prostaglandin E2 (3µM), rhWNT3a (60ng/µl), human r-spondin1 (250ng/ml), noggin (25 ng/ml), FGF10 (100ng/ml), human EGF (50ng/ml), human gastrin I (10nM), Y-27632 dihydrochloride (10µM), nicotinamide (10mM - Sigma Aldrich) and N-acetyl-L-cysteine (1mM - Sigma-Aldrich). For maintaining the PPDO, rhWNT3a, human r-spondin1 and noggin were substituted with the 3dGRO L-WRN conditioned medium supplement (Sigma-Aldrich - SCM105). Cells were plated using the Cultrex RGF basement membrane extract Type 2 (Bio-techne) as the matrix substance diluted 1:3 ratio with what we refer to as complete medium. Differentiation medium that was used for differentiation of organoids consisted of basal medium supplemented with N2 supplement (1X – ThermoFischer Scientific), B27-vitamin A supplement (1X – Life Technologies), A83-01 (5µM – Tocris Biosciences), Y-27632 dihydrochloride (10µM), nicotinamide (10mM-Sigma Aldrich) and N-acetyl-L-cysteine (1mM- Sigma-Aldrich). Screening medium used for the chemical screening experiments constituted of basal medium supplemented with: N2 supplement (1X – ThermoFischer Scientific), B27-vitamin A supplement (1X – Life Technologies), A83- 01 (5µM – Tocris Biosciences), prostaglandin E2 (3µM), FGF10 (100ng/ml), human EGF (50ng/ml), human gastrin I (10nM), Y-27632 dihydrochloride (10µM), nicotinamide (10mM-Sigma Aldrich) and N-acetyl-L-cysteine (1mM- Sigma-Aldrich).

Subculturing PPDO and HPDO was performed 7-10 days following previous splitting. Briefly, the Cultrex domes were dissociated with cold basal medium and organoids, washed 1X with basal medium and organoids were initially broken down manually with the P200 pipette. Following manual dissociation, organoids were incubated with TrypLE express (GIBCO) for 4 min at 37°C to further break the structures into smaller clamps and single cells and organoids were plated at the desired density with fresh cultrex and medium. All brightfield images across the manuscript were acquired with an EVOS M5000 cell imaging system with the 4X objective in brightfield mode, unless otherwise stated.

For differentiation test of PPDO, three different medium compositions were tested: 1- Basal organoid medium supplemented with 20% pig serum (Sigma-Aldrich). 2- Basal organoid medium supplemented with N2 supplement (1X – ThermoFischer Scientific), B27-vitamin A supplement (1X – Life Technologies), A83- 01 (5µM – Tocris Biosciences), Y-27632 dihydrochloride (10µM), nicotinamide (10mM-Sigma Aldrich) and N-acetyl-L-cysteine (1mM- Sigma-Aldrich) further supplemented with DAPT (1 µM - Tocris Biosciences), 4-Diethylaminobenzaldehyde (DEAB) (10 µM - Sigma-Aldrich) with/without BMS-536924 (5 µM – Sigma-Aldrich). For this medium combination, treatments performed either in Cultrex cultured organoids, or in dispersed cells cultured in mouse laminin (Corning laminin Sigma-Aldrich) coated ibidi chambers at 270µg/ml final concentration. 3- S5 and S6 media supplemented in PPDOs were prepared as described previously^75^.

### Human pancreatic ductal organoid culture

Exocrine pancreatic extracts were obtained following human islet isolation process from the Alberta Islet Distribution Program and the work was conducted under approval of the Health Research Ethics Board at the University of Alberta (Pro-00001620). Work with primary human tissue and HPDO generation and processing was performed under the ethical permit 2022-637-S-KH provided by the TUM ethics committee. HPDOs were generated either by handpicking ducts when available (death due to acinar enzyme tissue degradation) or by growing the single cell suspension from the byproduct of human islet isolation methods enriched in exocrine tissue^76^. HPDOs were passaged at least three times before any experimentation. Complete and screening media as well as passaging methodology used were identical to the PPDO cultures stated above.

### Immunostaining of PPDO

PPDO or HPDO were released from cultrex and washed once with cold PBS 1X followed by incubation with the cultrex organoid harvesting solution (Bio-Techne) for 30min. Organoids were fixed with 4% PFA for 45 min at 4°C, followed by incubation with PBS 1X supplemented with 0.1% Tween20 (Sigma-Aldrich). Cells were blocked in a solution of 0.2% w/v BSA (Sigma-Aldrich), 0.1% Triton X-100 (Sigma-Aldrich) in PBS1X for 1h at 4°C. Primary antibodies were incubated overnight in blocking solution at 4°C. Organoids were washed 2X for at least 1hr each in blocking solution and secondary antibodies were added overnight at 4°C together with DAPI to counterstain the nuclei. Following 2 washes for at least 1hr each at 4°C to remove unspecific binding of secondary antibodies, organoids were mounted with elvanol in 8-well chambers and imaged using a Leica SP5 or a Zeiss 880 laser scanning confocal microscope. Primary antibodies used in this study were: a-SOX9 (rabbit, 1:800, Millipore AB3555), a-KRT5 (rabbit, 1:200, abcam ab53121), a-KRT7 (mouse, 1:200, Agilent Technologies M701829-2), a-KRT8/18 (guinea pig, 1:1000, Origene BP5007), a-PAN-KRT (rabbit, 1:500, Agilent Z0622), a-BMPR1A (mouse, 1:400, LSBio LS-C191759), a-CDH1 (mouse, 1:800, BD Biosciences 610181), a-CDH1 (rat, 1:500, Takara M108), a-CFTR (rabbit, 1:100, Cell Signaling 78335), a-GP2 (mouse, 1:100, MBL D277-3), a-PDX1 (goat, 1:200, R&D systems AF2419), a-GCG (mouse, 1:200, Sigma-Aldrich G2654), a-AGR2 (rabbit, 1:200, Cell Signaling 13062S), a-GCG (guinea pig, 1:800, Takara M182), a-NKX6-1 (mouse, 1:100, Developmental Studies Hybridoma Bank F55A10), a-NEUROG3 (sheep, R&D systems AF3444), a- Pancreatic amylase (rabbit, 1:200, abcam 21156), a-INS (guinea pig, 1:200, LSBio (BIOZOL), LS-C85862-1), a-GCG (guinea pig, 1:1000, Takara, M182), Phalloidin Alexa Fluor 546 (Invitrogen A22283).

### Immunostaining of porcine pancreas

Cryosections (12 µm) of porcine pancreas were rehydrated in PBS 1X followed by permeabilization (0.3% Triton X-100), blocking (BSA-donkey serum-in PBS 1X) and overnight incubation with primary antibodies at 4°C. After washing primary antibodies, tissue was incubated with secondary antibodies plus DAPI to counterstain the nucleus for 2h at room temperature followed by washes and mounting with elvanol and 1.5 coverglass thickness. Images were obtained with a TCS SP5 confocal microscope. Primary antibodies against KRT7 (mouse, 1:200, Agilent Technologies M701829-2) and AGR2 (rabbit, 1:200, Cell Signaling 13062S) were used.

### CFTR assay

PPDOs were plated following standard splitting procedure mentioned above in 8-well glass bottom chambers (Ibidi). Following 4-5 days of culture, PPDOs were incubated with forskolin (10µM final concentration), secretin (1µg/ml final concentration) and a DMSO control and were imaged live using a Zeiss Observer microscope using brightfield lamp. Images from 5 independent positions/treatment were captured every 15min over the span of 4h. Lumen area was segmented with a custom-made model using the cellpose 2.0 algorithm^77^ and ROI area was calculated with Fiji^78^ for the first and last images of the acquisition.

### ScRNA-Seq

For the primary porcine pancreas dataset, pancreata from two pigs (WT and *INS*- eGFP) aged at PN150 were processed separately at two different days. Pancreata were cleaned from excess fat and connective tissue, chopped into fine pieces with scissors and incubated with 2mg/ml of collagenase V for 10min at 37°C. For the first pancreas, we manually isolated ductal fragments to enrich for the ductal cell population of the pancreas. Both isolated ducts and whole lysates from the second pancreas were incubated for 10 min with TrypLE Express (Gibco) to generate the single cell suspension. Single cell suspension was stained with 7-AAD (Invitrogen) as a live/dead cell marker. 20000 alive cells were sorted for sample #1. Sample #2 which contained the *INS*-eGFP transgene was stained with a conjugated Dolichos Biflorus Agglutinin (DBA) -biotin (1:100, Vector Laboratories B-1035-5) followed by incubation with Streptavidin Alexa Fluor 647 (1:500, Life Technologies S21374) to label ductal cells. Then 10000 DBA^+^ cells together with 10000 *INS*-eGFP cells and 20000 cells of the whole cell suspension were sorted using the FACSAria III (BD Biosciences) and all cells were pooled together.

For the organoid cultures, one confluent well of a 24-well plate containing PPDOs or HPDOs were detached from the cultrex using cold basal medium. The cells were washed once using cold basal medium and were incubated for 30 min in cultrex organoid harvesting solution to dissolve the extracellular matrix. PPDOs and HPDOs were dissociated to single cell suspension with TrypLE Express incubation for 10min at 37°C, followed by inactivation with a solution containing 1%BSA in PBS 1X. Cells were stained with 7-AAD (Invitrogen) as a live cell marker and 20000 live single cells were sorted using the FACSAria III (BD Biosciences).

The cell suspension was immediately used for single-cell RNA-seq library preparation with a target recovery of 10000 cells. Libraries were prepared using the Chromium Single Cell 3ʹ Reagent Kits v3.1 (10x Genomics, 1000268) according to the manufacturer’s instructions. Libraries were pooled and sequenced on an Illumina NovaSeq6000 with a target read depth of 50000 reads/cell. FASTQ files were aligned to the GRCh38 human genome with Ensembl release 111 annotations or to the in-house improved pig genome annotation^79^ and pre-processed using the CellRanger software v7.1.0 (10x Genomics) for downstream analyses.

### scRNA-Seq analysis

For downstream analysis, ambient RNA correction was performed using the SoupX tool^80^ using the autoEstCont function, except for the HPDO sample which was not deemed necessary to perform the correction. The corrected matrix was used for all the scRNA-Seq dataset analysis that was done with Scanpy^81^ suit of tools. For the PPDOs we filtered away genes expressed in less than 3 cells, cells with more than 3% mitochondrial gene counts, and cells containing more than 9000 and less than 600 (PN100) or 1000 (PN240). Counts were normalized to 10000 and log transformed using natural log and pseudocount 1, top 2000 highly variable genes were calculated using the ‘cell ranger’ flavour in the Scanpy package and clustering was performed using the top 20 PCs of the PCA with the Leiden algorithm with a resolution of 0.3 and 0.5 for the PN100 and PN240 samples respectively. Marker genes per cluster were identified using the Wilcoxon test and cell types/states were manually annotated based on marker genes and GO enrichment terms using the PANTHER^82^ database.

For the HPDO sample we filtered away genes expressed in less than 3 cells, cells with more than 15% mitochondrial gene counts, and cells containing more than 9000 and less than 1000 genes and 5000 counts. Counts were normalized to 10000 and log transformed using natural log and pseudocount 1, top 2000 highly variable genes were calculated using the ‘cell ranger’ flavour in the Scanpy package and clustering was performed using the top 25 PCs of the PCA with the leiden clustering algorithm with a resolution 0.5. Marker genes per cluster were identified using the Wilcoxon test and cell types/states were manually annotated based on marker genes and GO enrichment terms using the PANTHER^82^ database.

The two primary pancreas datasets were filtered separately before merging using 8% mitochondrial gene expression cutoff for both datasets followed by including cells for sample #1: with more than 400 and less than 9000 genes and for sample #2: with more than 800 and less than 9000 genes and more than 3000 UMI counts. For both samples doublets were identified and excluded using the scrublet^83^ tool using a doublet threshold of 0.25 and 0.15 respectively.

For integration PPDO and primary porcine datasets, we tested just aggregating the matrix, and integration using the scanorama^84^, scVI^85^ and sysVI^86^ tools. The scanorama and sysVI integration results are shown in the manuscript.

Ligand receptor interactions were inferred using the LIANA^87^ framework using the rank_aggregate function. Pig orthologues were used based on the human reference list using the built-in get_hcop_orthologs function from the HCOP database.

For mapping the HPDO to the tabula sapiens pancreas dataset^35^, we used the web interface of the scArches^88^ (https://www.archmap.bio/#/sequencer/genemapper) to upload our filtered HPDO scRNA-Seq matrix and used the scVI^85^ and kNN approach to map our HPDO dataset to the atlas. Visualization using the umap was performed using the Scanpy suit.

### RNA-Seq

For the bulk RNA-Seq analysis, PPDOs cultured in complete or differentiation media were collected after 7 days of culture and washed twice with PBS 1X. RNA was extracted using the Arcturus PicoPure RNA isolation kit (Applied Biosystems). RNA quality was assessed using the bioanalyzer and all samples showed a RIN value more than 9 and therefore included in library preparation using the mRNA prep ligation kit (Illumina) with poly-A tail selection, following the kit’s instructions. The libraries were sequenced using the NovaseqX+ sequencer (Illumina) with a depth of ≥ 30 Million reads per sample (paired-end mode 2×100 bases). Library preparation and sequencing were performed at Helmholtz Zentrum München (HMGU) by the Genomics Core Facility. Processing of the sequencing that generated the aligned reads was performed using the nf-core pipeline (v3.14) with Nextflow (23.10.1) and the salmon tool used for quantification of transcripts^89–95^. The reads were aligned to the custom annotated Sus Scrofa genome.

The gene length scaled counts output file from salmon was used for differential expression analysis. Raw counts from transcripts with –iso and –ext were aggregated to get the summed counts per gene transcript. The DESeq2^96^ package was used for differential gene expression analysis with the apeglm^97^ log fold change shrinkage applied for effect size shrinkage and the EnhancedVolcano package for visualization. GO enrichment analysis was performed with the PantherDB^82^ database.

### Chemical screen

PPDOs/HPDOs were dissociated into single cells as described above for passaging and 7µl droplets were plated in a glass-bottom, 96-well plates. We cultured PPDOs/HPDOs for 2 days in complete media for organoid formation. After 2 days, cultures were washed once with PBS 1X, and chemicals from a Tocris-FDA approved drugs library (Tocris-7200) were added in a final concentration of 20µM in medium without WNT3a-Rspondin-noggin factors. 4 control wells/plate were used with complete and media without the 3 factors supplemented with DMSO (Sigma-Aldrich). PPDOs were incubated with the drugs for 3 days, followed by centrifugation of the plate for 10’, at 900g at 10°C and fixation with 4% paraformaldehyde. PPDOs/HPDOs were permeabilized with 0.5% Triton X-100 solution for (1hr-room temperature), followed by blocking (1hr-room temperature) with 3% donkey serum supplemented with 0.1% Triton X-100 blocking buffer. PPDOs/HPDOs were incubated with pHH3 (1:1000, Cell Signaling) primary antibody overnight at 4 °C followed by secondary antibody incubation together with DAPI to counterstain nuclei (2h-room temperature). PPDOs/HPDOs were mounted with Vectashield (Vector Laboratories) and the whole 96-well plate was imaged using a Cell Discoverer 7 microscope (Zeiss) using the laser-scanning confocal mode and imaging 2 random positions/well. The whole plate was also imaged using the brightfield mode to assess the survival of the cultures. To count the percentage of proliferating cells both the DAPI and pHH3 channels were segmented in 3D using the Cellpose 2.0 algorithm^77^ on custom-trained models (trained on the 2D images of the experiment), the proliferation percentage calculated as the number of pHH3^+^ to total DAPI^+^ cells and the z-score was calculated in reference to complete media using the mean and standard deviation measures. The counts from the two positions/well imaged were averaged to get an overall proliferation percentage/well. For the secondary screen, all chemicals were purchased from different vendors and diluted fresh to avoid any confounding effects using the same batch of drugs and focus on the most promising biological data. The following chemicals were purchased and screened at a final concentration of 20µM: losartan potassium (Sigma-Aldrich), telmisartan (Santa Cruz Biotechnology), cimetidine (Sigma-Aldrich), famotidine (Sigma-Aldrich), tranylcypromine hydrochloride (Cayman Chemicals), GSK-LSD1 hydrochloride (Tocris Biosciences), picrotoxin (Sigma-Aldrich), flumazenil (Cayman Chemicals).

